# A rationally designed neuraminidase immunogen elicits humoral responses to a conserved viral site

**DOI:** 10.64898/2026.08.08.743685

**Authors:** Rochel Hecht, Faez Amokrane Nait Mohamed, Shiyu Zhang, Shaun Rawson, Saeyoung E. Lee, Connor L. Murphy, Dana Thornlow Lamson, Larance Ronsard, Timothy M. Caradonna, Daniel Lingwood, Aaron G. Schmidt

## Abstract

Efforts to develop a universal influenza vaccine have primarily focused on the surface-exposed viral hemagglutinin (HA), but neuraminidase (NA) is an additional target for cross-reactive, protective responses. Here, we used hyperglycosylation as an immunogen design approach to reshape anti-NA humoral immunity toward conserved antigenic regions. Iterative design produced a hyperglycosylated NA immunogen that retained enzymatic activity and reactivity to a conformation-specific antibody recognizing the conserved catalytic site. In mice, the hyperglycosylated immunogen elicited serum antibody responses of comparable magnitude to those elicited by the wild-type NA immunogen. However, serum antibodies elicited by the hyperglycosylated immunogen had increased breadth, recognizing N2 NAs from H2N2 and H3N2 viruses spanning nearly 65 years of antigenic drift, as well as a heterosubtypic N9 NA. Single B cell analyses identified a monoclonal antibody that competed with a component of the serum antibody response elicited by the hyperglycosylated immunogen, and structural characterization showed that it recognizes a previously undefined, conserved epitope at the NA tetramer interface. Passive transfer of this interface-directed antibody partially protected mice against lethal heterologous influenza challenge. Collectively, these data show that glycan shielding can reshape and enrich humoral responses toward a conserved antigenic region on NA. The immunogen and hyperglycosylation design strategy described here provide a template for developing next-generation NA-based influenza vaccines.

**ONE SENTENCE SUMMARY:** A hyperglycosylated influenza neuraminidase immunogen reshapes humoral immunity and elicits antibodies targeting a conserved tetramer-interface epitope.

## INTRODUCTION

Influenza is an annual public health burden, and its mutational capacity raises the threat of future pandemics^1^. Its two surface proteins, hemagglutinin (HA) and neuraminidase (NA), are both subject to antigenic drift under pressure from population humoral immunity. While most immunogen design efforts have concentrated on HA, NA is also the target of protective antibodies. NA mutates discordantly and less rapidly than HA^2,3^ and, in humans, anti-NA antibody responses are an independent correlate of protection against influenza infection^4^. Antibodies that target NA can be broadly reactive, recognizing different strains and, in some cases, multiple NA subtypes. The broadly reactive antibodies often engage conserved sites on NA, including the catalytic site (CS)^5–8^ and the recently defined NA underside^9,10^. Although NA is present in some seasonal influenza vaccines, its antigen content is not standardized, contributing to variability in NA quantity, quality, and immunogenicity across vaccine formulations^11,12^. However, eliciting broad NA antibodies through inclusion of an NA component in next-generation influenza vaccines will likely augment vaccine effectiveness and protect against future pandemics^13–15^. Indeed, previously established anti-NA immunity was thought to contribute to the relative protection of vulnerable populations during the 1968 H3N2 and 2009 H1N1 influenza pandemics^16–19^. Although several conserved NA epitopes have been structurally defined, relatively few are known compared with HA, and additional conserved antigenic site(s) could both inform vaccine design and uncover new mechanism(s) of antiviral protection.

A major goal of a next-generation influenza vaccines has been to elicit humoral immunity against conserved sites on HA ^20^ including the receptor binding site (RBS)^21,22^, stem^23,24^, and the interface epitope^25–27^; it has also been proposed for conserved sites on NA, including the CS and underside^28,29^. However, these antigenic regions are generally immunologically subdominant following infection or vaccination. Accordingly, rational immunogen design strategies have been developed to alter immunodominance and redirect humoral responses toward conserved epitope(s). One approach is to introduce non-native glycans onto the protein surface, resulting in hyperglycosylated immunogens. The introduced glycans sterically occlude selected antigenic regions, reducing accessibility to B-cell receptors and antibodies while preserving exposure of desired epitope(s). This approach is predicated on the naturally varying glycosylation patterns on surface-exposed viral proteins, such as the HIV envelope protein, HA, and NA that contribute to evasion of host humoral immunity^30–32^. Indeed, varied glycosylation patterns on HA influence its overall antigenicity and can contribute to reduced vaccine effectiveness^30,33–35^.

Previously, we showed that hyperglycosylating HA can modulate elicited humoral immunity without affecting overall antigenicity^25,36^. Additionally, these hyperglycosylated HA immunogens focused the humoral response and defined the conserved trimer interface epitope as a target of humoral immunity that is also observed in humans^25–27^. Here, we applied hyperglycosylation to influenza NA to determine whether glycan-mediated immune focusing could redirect antibody responses toward conserved NA epitopes while preserving immunogenicity. We designed a hyperglycosylated NA immunogen based on the historical H2N2 NA from1957, aiming to redirect humoral immunity with a particular emphasis on focusing to the CS. The biochemically and biophysically characterized hyperglycosylated NA immunogen retained its immunogenicity relative to the wild-type NA. Subsequent sera and single B cell analyses showed that this immunogen focused humoral immunity with enhanced N2 NA breadth. An isolated monoclonal antibody recognized an occluded epitope at the NA tetramer interface, similar to the HA trimer interface-epitope described previously. Lastly, this NA interface-directed antibody is partially protective against a lethal heterologous influenza challenge. Our results show that hyperglycosylating NA can influence humoral responses and subsequently focus the humoral immune response to a previously undefined, conserved epitope. Incorporating this antigenic region into immunogens for next-generation influenza vaccines may contribute to broadly protective NA-based immune responses.

## RESULTS

### Design and characterization of neuraminidase immunogens

Neuraminidase (NA) has 11 subtypes and 2 lineages within the influenza A and B viruses, respectively (**Fig. 1A**) ^37^. It is a surface-exposed, homo-tetrameric glycoprotein with a stalk leading into a transmembrane domain and cytoplasmic tail (**Fig. 1B**)^38^. The catalytic site (CS) at the center of each beta-propeller “head” domain is distal to the virion membrane and is conserved across subtypes (**Fig. 1B and fig. S1A, B**)^37^. We selected the historical H2N2 A/Japan/305/1957 (J’57) NA as the prototypic antigen for our immunogen design efforts because this influenza strain is the earliest human N2 lineage preceding decades of H3N2 antigenic drift and therefore provides a useful template for evaluating subsequent breadth of elicited responses. J’57 NA has 5 native predicted N-linked glycosylation sites (PNGs) per monomer (**Fig. 2A**). To increase the number of PNGs we identified 10 additional historical PNG sites that naturally occurred during influenza evolution by analyzing ∼7,500 unique HxN2 strains. After modeling the native and historical PNGs we introduced an additional novel PNG site (Asn336) to occlude the remaining solvent accessible NA surface while leaving the CS site accessible (**Fig. 2B and fig. S2A**).

**Fig. 1:**
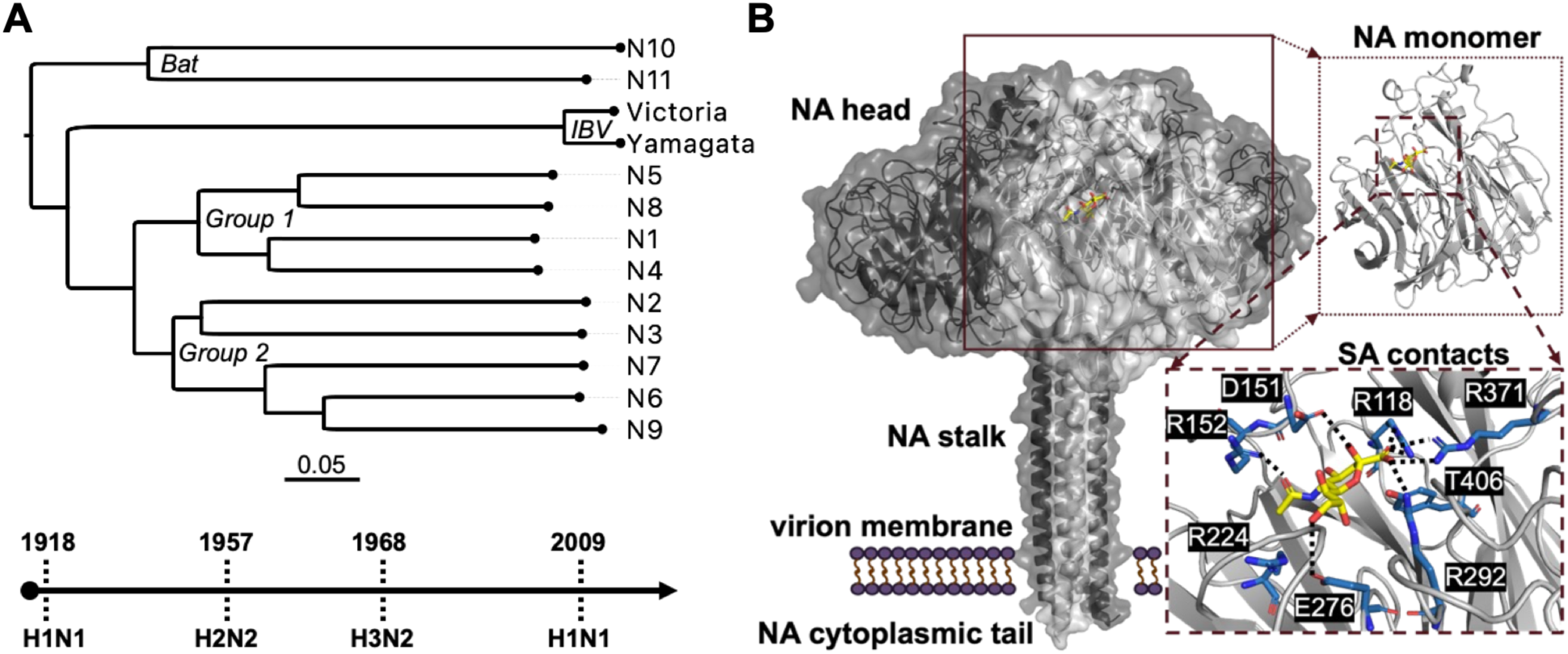
Neuraminidase (NA) subtypes and conserved features. **(A**) NA phylogenetic tree with 9 IAV subtypes, two IBV lineages, and two bat-associated subtypes. Below, timeline with four global pandemics with their associated IAV subtypes. **(B)** Ribbon and surface representation of NA (PDB 6Q20^16^), with its homo-tetrameric head, stalk, transmembrane domain, and cytoplasmic tail indicated. Conserved residues of the catalytic site (CS) that contact sialic acid (inset) are marked (see also **fig. S1A-B**). The NA stalk is represented by the non-native bacterial tetrabrachion (TB) domain (PDB 1FE6^41^). One NA monomer is shown in light gray for clarity.

**Fig. 2.**
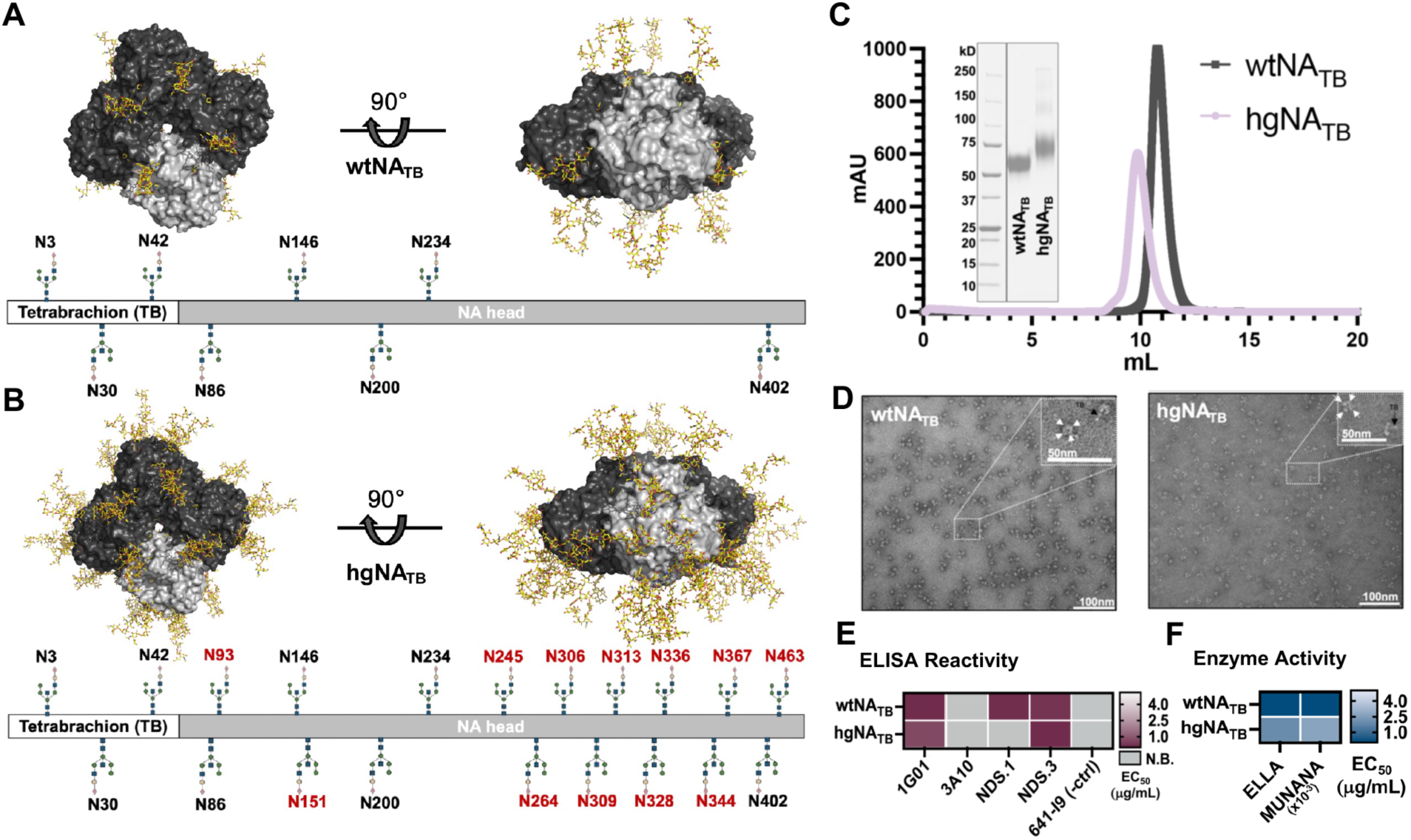
Design and characterization of wildtype and hyperglycosylated NA proteins. **(A)** Wildtype J’57 N2 NA tetramer (wtNA_TB_) shown in surface representation with 5 native PNGs modeled. Schematic representation showing the N-terminal tetrabrachion (TB) domain and native glycan positions (using N2 amino acid numbering). The PNG at position N30 on TB is non-native. **(B)** Hyperglycosylated J’57 N2 NA tetramer (hgNA_TB_) with 11 additional glycans modeled; schematic representation shown below includes non-native glycans in red. **(C)** Size-exclusion chromatogram of the wtNA_TB_ and hgNA_TB_ proteins and SDS-PAGE analysis. **(D)** Representative negative stain electron micrographs of wtNA_TB_ and hgNA_TB_ proteins. White and black arrows indicate individual NA monomers (∼5nm wide) and the TB domain, respectively. **(E)** Reactivity of CS-directed mAbs, 1G01, 3A10, and underside-directed mAbs, NDS.1 and NDS.3, against wtNA_TB_ and hgNA_TB_ proteins in ELISA. EC_50_ (µg/ml) are listed. **(F)** Enzymatic activity of wtNA_TB_ and hgNA_TB_ proteins in ELLA (large fetuin substrate) and MUNANA (small fluorescent substrate) assays.

To enhance recombinant expression and tetramer formation, we replaced the native NA stalk with the bacterial tetrabrachion domain (TB)^39^ and introduced a PNG site to mitigate its potential immunogenicity (**Fig. 2A**,**B****, and fig. S2A**). The wildtype NA (wtNA_TB_) and hyperglycosylated NA (hgNA_TB_) were recombinantly expressed from mammalian cells to ensure complex glycosylation^40^. Proteins were purified to homogeneity and monodispersity and visualized using negative stain electron microscopy (**Fig. 2C,D**). Both proteins migrated similarly on SDS PAGE after deglycosylation using PNGase (**fig. S2B,C**). The wtNA_TB_ and hgNA_TB_ proteins had comparable reactivity to the conformation-specific CS-directed 1G01 monoclonal antibody (mAb), and each had enzymatic activity in both ELLA and MUNANA assays (**Fig. 2E,F**). These data show that extensive NA hyperglycosylation did not alter proper folding, recognition of the CS by a conformation-specific antibody, and enzymatic activity.

### Immunogenicity of neuraminidase immunogens

We next assessed the immunogenicity of the wtNA_TB_ and hgNA_TB_ proteins in C57BL/6 mice using a homologous prime-boost-boost regimen (**Fig. 3A**). While hgNA_TB_-immunized sera reacted comparably to both the hgNA_TB_ and wtNA_TB_ proteins, wtNA_TB_-immunized sera had significantly reduced reactivity to the hgNA_TB_ protein (**Fig. 3B,C** **and fig. S3A**). Thus, extensive NA hyperglycosylation did not measurably reduce overall serum immunogenicity and could occlude antigenic site(s) recognized by the wtNA. Additionally, compared with sera elicited by wtNA_TB_ immunizations, hgNA_TB_-immunized sera had greater breadth, recognizing the NA from a contemporary H3N2 strain, A/Darwin/6/2021, and a heterosubtypic H11N9 strain, A/tern/Australia/G70C/1975 (**Fig. 3D and fig. S3B**). To assess whether this enhanced breadth was a consequence of serum antibody focusing to the CS, we introduced a glycan into the CS of the wtNA_TB_ (ΔCS) that abrogated binding to CS-directed 1G01 and 1E01 mAbs (**fig. S3C,D**). If the enhanced breadth resulted primarily from CS-directed antibodies, we would expect substantially reduced serum binding to the ΔCS construct. While hgNA_TB_-immunized sera had a slight reduction in reactivity to the ΔCS construct relative to wtNA_TB_-immunized sera (**Fig. 3E,F**) sera from both cohorts inhibited NA enzymatic activity comparably (**Fig. 3G**). These findings suggest that serum antibodies recognizing epitope(s) outside the CS contributed substantially to the enhanced breadth observed following hgNA_TB_ immunization. There was no appreciable difference in sera reactivity to monomeric versus tetrameric versions of the immunogens, suggesting that a minimal proportion of sera antibodies were likely targeting complex epitopes spanning NA protomers (**fig. S3E**). Lastly, to determine the extent of tag-specific responses, we appended the unrelated HA “head” domain from H1 Solomon Isalnds/03/2006 onto the TB tetramerization domain and assayed sera from both the wtNA and hgNA cohorts. Despite the introduced glycan on the TB domain, both cohorts had detectable tag-directed responses (**fig. S3F**); however, these responses did not appear to reduce NA-specific serum responses. Collectively, these data show that both wtNA_TB_ and hgNA_TB_ were immunogenic and that the enhanced breadth elicited by hgNA_TB_ was more consistent with responses targeting an epitope(s) outside the CS than with enhanced CS focusing.

**Fig. 3.**
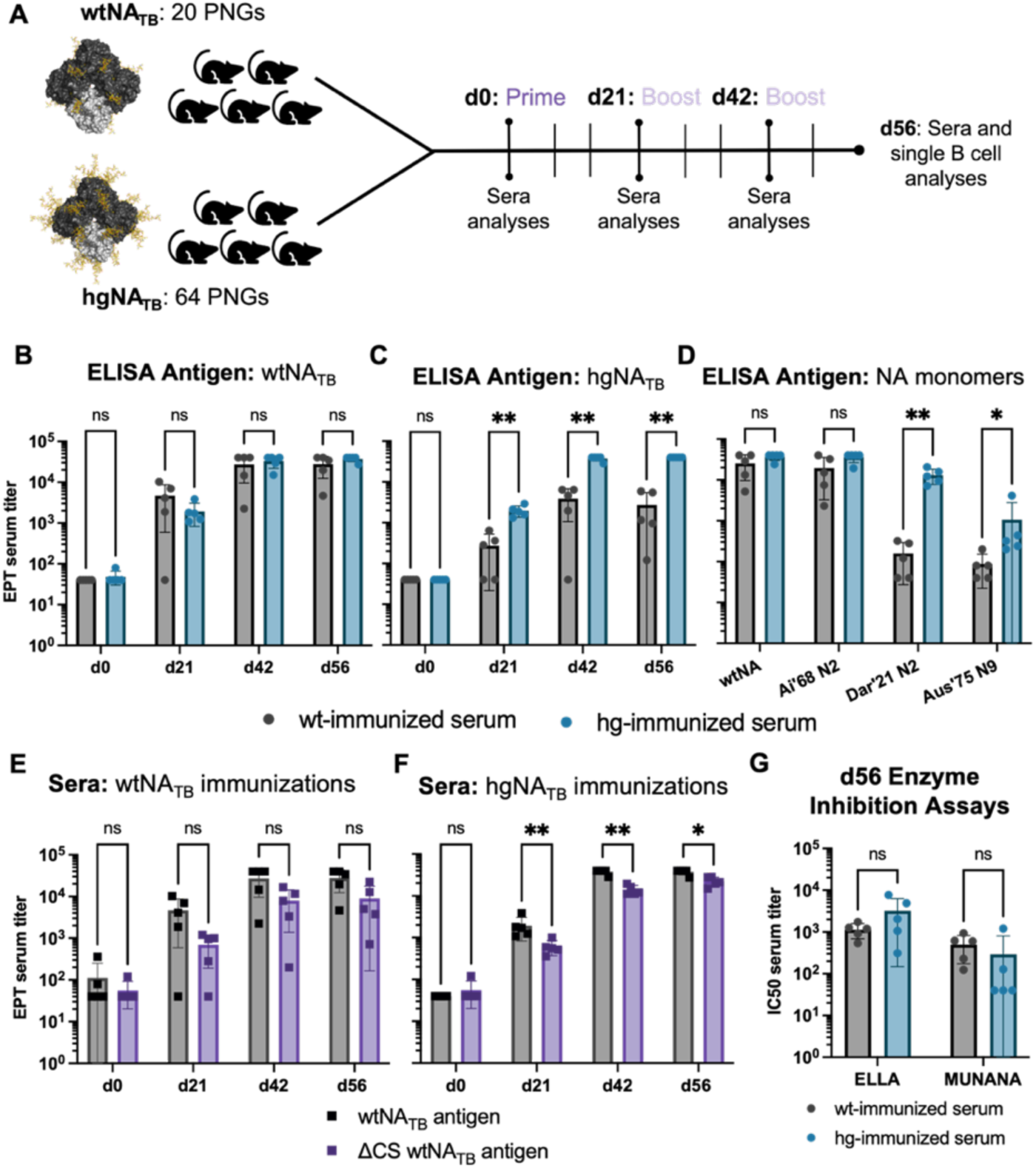
***In vivo* testing and serum characterization. (A)** Immunization regimen and downstream analyses for wtNA_TB_ and hgNA_TB_ immunogens. Sera reactivity from hgNA_TB_ and wtNA_TB_ immunizations to the **(B)** wtNA_TB_ and **(C)** hgNA_TB_ antigens. **(D)** Reactivity of d56 sera from hgNA_TB_ and wtNA_TB_ immunizations to contemporary monomeric N2 and N9 NAs. Sera reactivity to the ΔCS wtNA antigen from **(E)** wtNA_TB_ and **(F)** hgNA_TB_ immunizations. **(G)** Enzymatic activity inhibition by d56 sera in ELLA and MUNANA assays. Reactivity data were compared using nonparametric, unpaired Mann-Whitney tests. Lines on graphs represent the mean and standard deviation. p-values: **0.008, *0.02.

### Single B cell analysis and reactivity profiles from neuraminidase immunogens

We next isolated single B cells elicited by the wtNA_TB_ and hgNA_TB_ immunogens to determine whether the altered serum breadth was reflected in the underlying B-cell repertoire. We also aimed to characterize the genetic features of elicited B cells and potentially isolate monoclonal antibodies (mAbs) that might explain the observed breadth of the hgNA_TB_-immunized sera. 188 B cells from two mice per cohort (n=376 B cells per cohort) were index-sorted based on their antigen-specific binding profiles (**fig. S4**). Consistent with the serum reactivity profiles, there were statistically fewer single B cells from wtNA_TB_ cohort that bound hgNA_TB_ (**Fig. 4A**). Biased gene usage was observed in both variable heavy (IGHV; V_H_) and kappa light (IGKV; V_K_) chain repertoires. IGHV1-85 and IGKV12-41 accounted for ∼45% and ∼83%, respectively, of the variable genes from hgNA_TB_-immunized mice but only ∼2% and ∼25%, respectively for the wtNA_TB_-immunized mice (**Fig. 4B,C** **and fig. S5A-D**). Complementarity-determining region (CDR) 3 of the heavy chain length was roughly equivalent, centered around ∼15 amino acids, and CDR L3 length was almost exclusively 13 amino acids (**fig. S5E,F**).

**Fig. 4.**
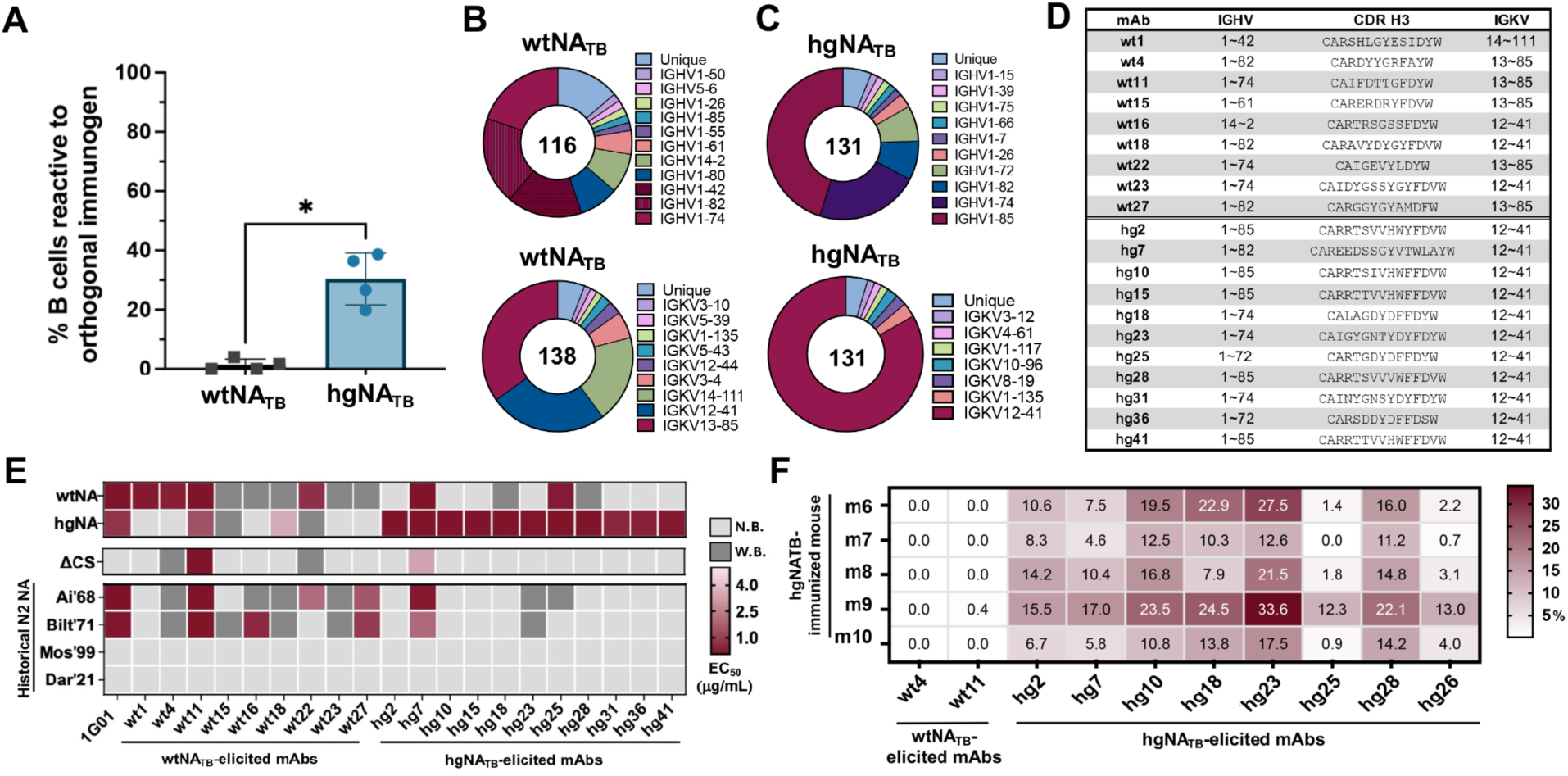
Single B cell gene usage and binding profile analyses. **(A)** Comparison of reactivity of isolated B cells from wtNA_TB_-immunized mice against the hgNA_TB_ protein and isolated B cells from hgNA_TB_- immunized mice against the wtNA_TB_ protein. Genetic analysis of isolated BCR **(B)** heavy and **(C)** light chains. **(D)** Representative recombinantly expressed mAbs from the wtNA_TB_- and hgNA_TB_ -immunized mice (wt and hg, respectively) variable heavy and light chain genes and CDR H3 amino acid sequences. **(E)** Reactivity of select mAbs to N2 NA proteins in ELISA. **(F)** hgNA_TB_-immunized sera competition with select mAbs. Data were compared using Mann-Whitney tests. p-value: *0.029. (W.B.= EC_50_ between 10- 50 μg/mL; N.B. = no binding).

Of the total indexed cells, we obtained 38 and 45 paired sequences from wtNA_TB_- and hgNA_TB_- immunized cohorts, respectively. Of these, 32 and 38 mAbs were successfully expressed (**Fig. 4D**). Approximately 40% of the wtNA_TB_-elicited mAbs were reactive against the hgNA_TB_ antigen, while ∼84% of the hgNA_TB_-elicited mAbs were reactive against the wtNA_TB_ antigen (**fig. S5G**), consistent with the serum and B cell binding profiles (**Fig. 3B,C** **and** **Fig. 4A**). We next downselected mAbs from each cohort based on overrepresented gene usage, CDR H3 identity, and/or distinctive binding profiles to assess breadth, potential epitope-focusing, and serum competition. Although some wtNA_TB_-elicited mAbs bound historical H3N2 between 1968-1971, none of the recombinant mAbs from either cohort bound contemporary N2 NAs after 1971. Indeed, the isolated hgNA_TB_-elicited mAbs were restricted in their binding to the hgNA_TB_ immunogen itself and, for a limited number of mAbs, to wildtype J’57 NA (**Fig. 4D**). wt1 and hg25 mAbs appeared sensitive to the ΔCS NA suggesting an epitope that partially overlaps with the CS. However, while some of the tested mAbs, including wt1 and hg25, inhibited NA catalytic activity in the ELLA assay (**fig. S6A**), none inhibited in the MUNANA assay (**fig. 6B**), suggesting that they did not overlap directly with the CS (like 1G01); the vast majority of the isolated wt- and hgNA_TB_-elicited mAbs therefore likely engage epitope(s) outside of the CS. A subset of hgNA_TB_- elicited mAbs and two wtNA_TB_- elicited mAbs were used for competition with the hgNA_TB_- immunized sera. Neither wt4 nor wt11 mAbs competed with sera from hgNA_TB_ immunizations. While hg10, 18, and 23 mAbs had modest inhibition with sera across each mouse, these mAbs did not bind Dar’21 NA and therefore could not be used to define the conserved N2 epitope(s) contributing to the observed sera breadth **(Fig. 4F**).

**Fig. 5.**
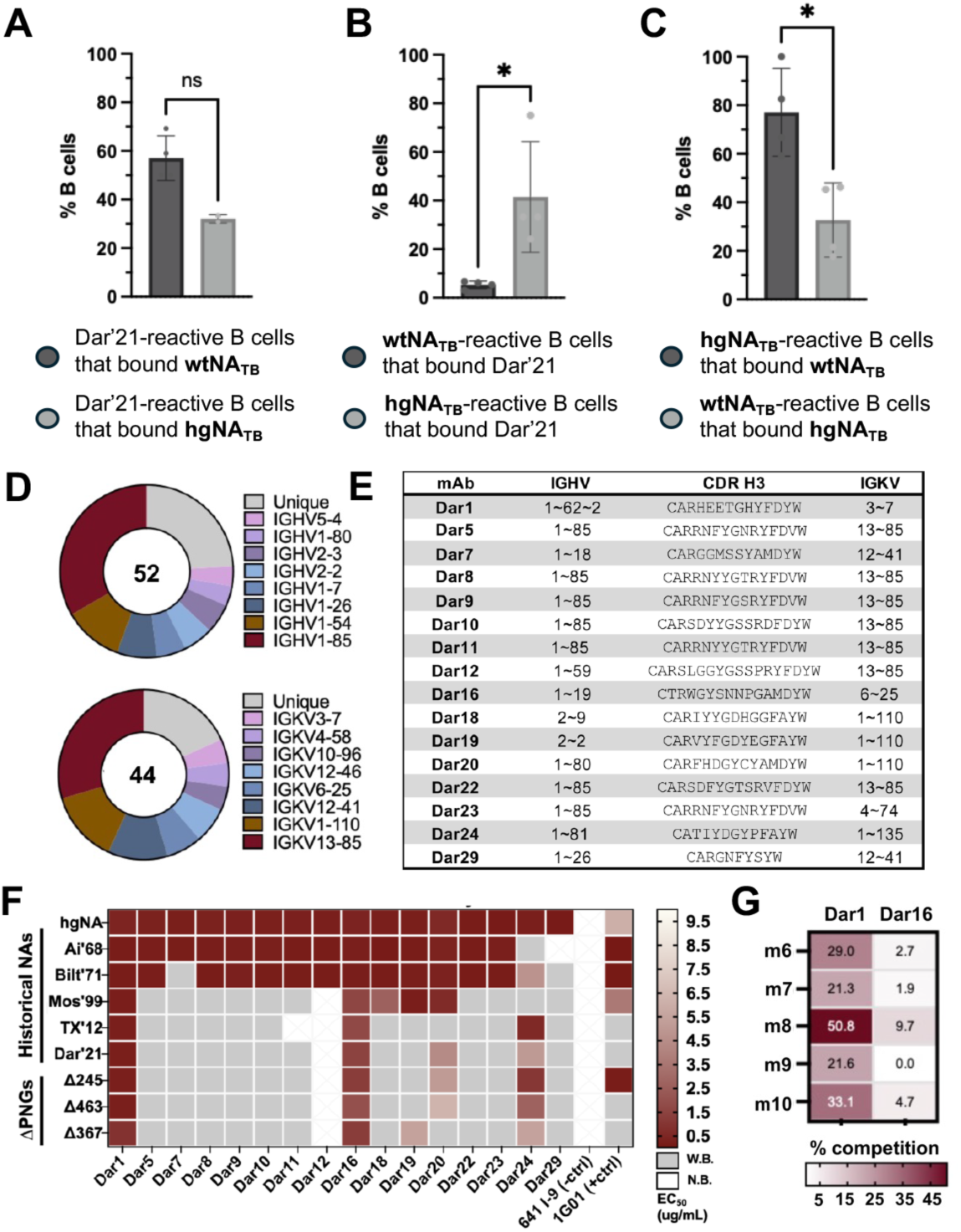
Single B cell gene usage and binding profile analyses from Dar’21 NA-reactive B cells. **(A)** Comparison of Dar’21 NA reactive B cells that also bound wtNA_TB_ and hgNA_TB_ proteins. **(B)** Comparison of hgNA_TB_- or wtNA_TB_-reactive B cells that also bound Dar’21 NA. **(C)** Percentage of hgNA_TB_-reactive B cells that also bound wtNA_TB_ protein compared with the percentage of wtNA_TB_-reactive B cells that also bound hgNA_TB_ protein. **(D)** Genetic analyses of B cells isolated from hgNA_TB_-immunized mice that bound Dar’21 NA. **(E)** Recombinantly expressed mAbs from Dar’21 NA-specific sort with their variable heavy and light chain genes and CDR H3 amino acid sequences. **(F)** Reactivity of isolated mAbs to N2 NAs. **(G)** Competition of Dar1 and Dar16 mAbs with sera from the hgNA_TB_-immunized cohort. Statistical analysis was performed using Mann-Whitney tests. Bars and lines represent mean and standard deviation, respectively. p-values: *0.029. (N.B. = no binding; W.B.= EC_50_ between 10-50 µg/mL).

**Fig. 6:**
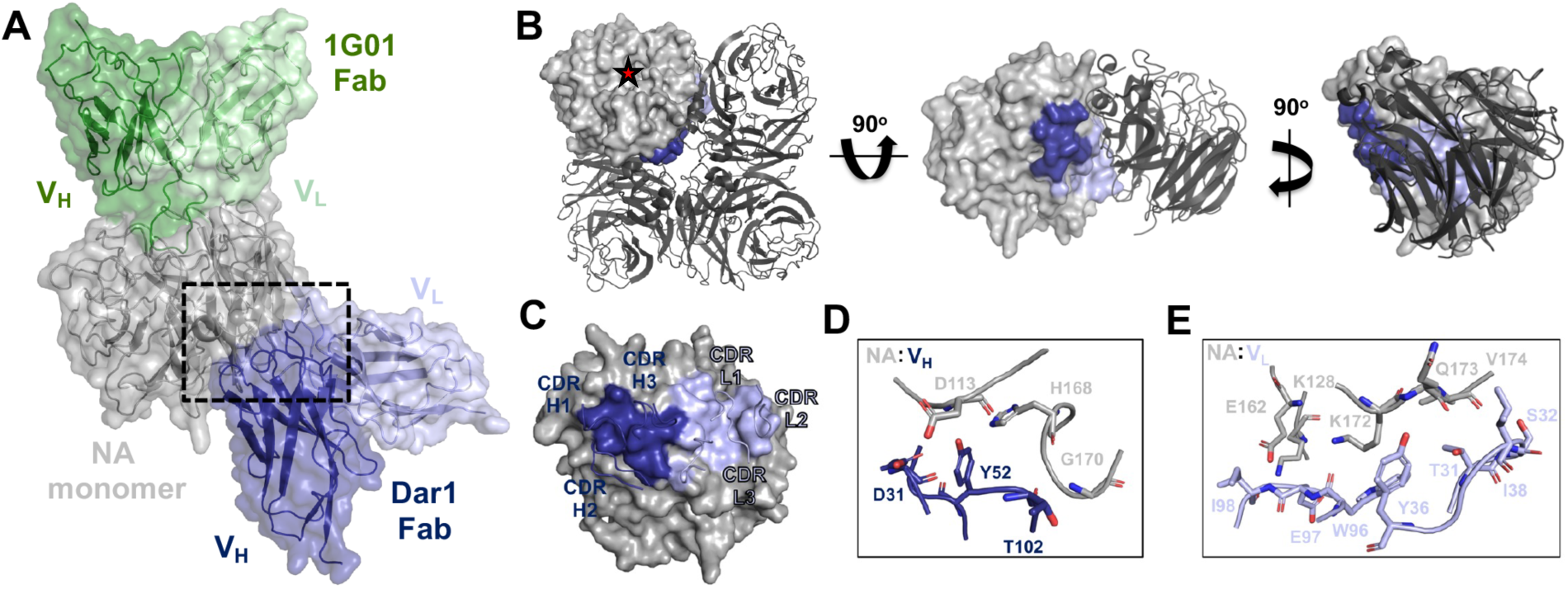
Structural characterization of Dar1 mAb. **(A)** Cryo-EM structure of Dar1 and 1G01 Fabs in complex with a J’57 NA monomer. Surface representation is shown with V_H_ and V_L_ domains of each Fab noted. Dashed rectangle marks the Dar1:NA interface illustrated in subsequent panels. **(B)** Mapping of the occluded Dar1 footprint on the NA tetramer (PDB 3TIA). An NA protomer with the Dar1 footprint is shown in surface representation in context with the NA tetramer. The initial orientation is of the NA tetramer perpendicular to the virion membrane with the catalytic site noted in one protomer (red star). Rotation and hiding of two NA protomers exposes the otherwise occluded V_H_ footprint. Final rotation shows how an adjacent NA protomer occludes the V_L_ footprint. **(C)** Dar1 footprint shown as colored surface representation on the NA monomer with CDRs from V_H_ (dark blue) and V_L_ (light blue). NA interactions with **(D**) V_H_ and **(E)** V_L_ residues in the antigen binding site.

### Isolation of single B cells reactive to contemporary N2 neuraminidase

Since the isolated mAbs did not fully explain the breadth observed in serum, we used the contemporary Dar’21 NA and isolated additional B cells to identify cross-reactive mAbs for serum competition studies and structural studies (**fig. S7A-D**). We obtained 52 heavy chain and 44 light chain sequences from hgNA_TB_-immunized mice (**Fig. 5A and fig. S8A-D**) and downselected 16 mAbs for recombinant expression based on gene usage and CDR H3 identity (**Fig. 5E**). 16 mAbs bound both hgNA_TB_ and wtNA_TB_ with high avidity (EC_50_ <1μg/mL), and many had broad but relatively weak (EC_50_ ∼10-50 μg/mL) reactivity to historical N2 NA strains from 1957-2021 (**Fig. 5F**). However, two mAbs, Dar1 and Dar16, bound tightly to all tested historical and contemporary NAs, as well as the hgNA_TB_ and wtNA_TB_ immunogens. We next designed three mutants on Dar’21 NA that abrogated the shared glycans between J’57, Dar’21, and the hgNA_TB_ immunogen, Δ245, Δ367, and Δ463, to assess glycan-dependent interactions; Dar1 and Dar16 were not critically dependent on any of these glycans (**Fig. 5B and fig. S9A**). Furthermore, both Dar1 and 16 did not inhibit NA activity in the ELLA assay, nor did Dar1 compete with mAbs engaging the conserved underside epitope (**fig. S9B,C**). Unlike Dar16 however, Dar1 competed substantially with sera from hgNA-immunized mice, suggesting its footprint overlapped with a component of the elicited polyclonal serum response (**Fig. 5G**).

### Structure of an elicited cross-reactive antibody

We determined a cryo-EM structure of Dar1 Fab in complex with J’57 NA monomer and the CS-directed 1G01 Fab to 2.8Å (**Fig. 6 and fig. S10**). Dar1 binds a region outside the CS at the tetramer interface between adjacent NA protomers (**Fig. 6A,B** **and fig. S11A**). The Dar1 epitope overlaps significantly with the interface created by a single adjacent NA monomer within the tetramer and is largely contributed by the V_L_ domain while the V_H_ domain has a footprint at the exposed four-fold axis. The total buried surface area (BSA) in the antigen combining site is equally distributed across both the V_H_ and V_L_ with ∼430 and 450 Å^2^, respectively, and is roughly equivalent to the ∼900Å^2^ contributed by the V_H_ alone of 1G01 within the CS (**Fig. 6C**). The few hydrogen bonding interactions between Dar1 and NA include the hydroxyl of Y36 in CDRL1 with the backbone of E162 on NA and the backbone of I98 in CDRL2 with Q173 on NA (**Fig. 6D,E**); these NA residues include key salt bridge and hydrogen bonding with an adjacent NA protomer in the NA tetramer. The Dar1 epitope is highly conserved across N2s and thus explains the observed breadth (**fig. S11B**). This contrasts with the CS antigenic region which is highly conserved only within the NA “core” with the surrounding periphery variable (**fig. S11B**). This variability contributes to reduced reactivity and breadth of CS-directed antibodies including 1G01--indeed, 1G01 does not recognize the contemporary Dar’21 NA while Dar1 does. The Dar1 epitope is distinct from any other previously described murine or human NA-directed antibodies and draws a parallel to antibodies recognizing the HA trimer interface^26,27^.

### Protection by Dar1 antibody against heterologous influenza challenge

We next assessed the protective activity of Dar1 against heterologous influenza challenge. C57BL/6 mice (n=10 per cohort, 5 male and 5 female) were administered Dar1, 1G01, or Z021 (Zika virus-specific, negative control) IgGs and then challenged intranasally with a lethal dose of X-31 H3N2 influenza virus. Mice receiving prophylactic 1G01 or Dar1 IgG followed similar recovery trajectories with 90% and 60% survival, respectively, whereas all mice receiving Z021 IgG were euthanized by day 6 (**Fig. 7A, B**). These data show that an antibody targeting a conserved epitope outside the CS can confer partial protection against lethal heterologous challenge and support further investigation of this antigenic region as a target for next-generation influenza vaccines.

**Fig. 7:**
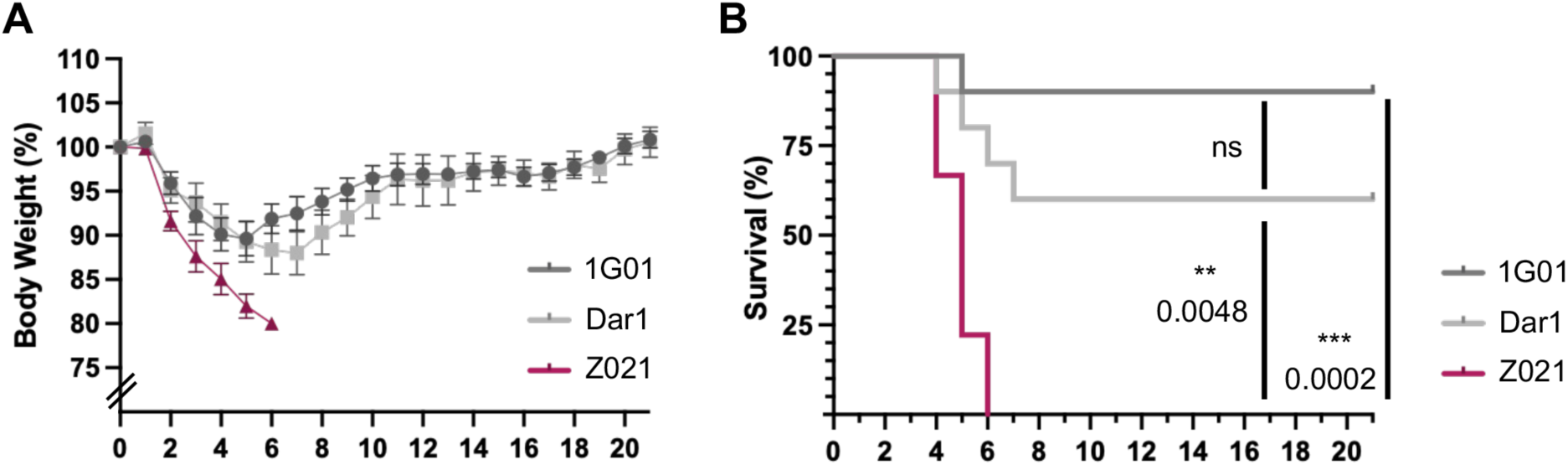
Protective efficacy of Dar1 mAb against heterologous H3N2 challenge. **(A)** Percent body weight of each mouse relative to their starting weight on day 0 (n=10 mice per tested antibody, 5 males and 5 females). **(B)** Survival curves of cohorts receiving Dar1, 1G01 (CS-directed, positive control), or Z021 (Zika virus-specific, negative control) IgGs after a lethal influenza infection. Lines and bars indicate mean weight and standard error, respectively. Data were analyzed using Gehan-Breslow-Wilcoxon tests. P-values are noted.

## DISCUSSION

Here, we showed that non-native glycans can be introduced onto a prototypical N2 NA while preserving enzymatic activity and reactivity to a conformation-specific antibody recognizing the conserved CS. The hyperglycosylated NA immunogen reshaped humoral immunity toward a previously uncharacterized conserved epitope at the tetramer interface that is uniquely conserved across the N2 subtype from its H2N2 introduction to humans in 1957 through more contemporary H3N2 strains in 2021. Serum competition showed Dar1-like responses targeting this antigenic region contributed to the polyclonal serum response elicited by the hyperglycosylated immunogen. Despite it being an occluded epitope, passive transfer of Dar1 mAb provided partial protection against a heterologous viral challenge, suggesting this epitope is sufficiently accessible *in vivo* to confer antibody-mediated protection. These data show how strategic placement of glycans on the NA surface can alter humoral responses and immune focus to a highly conserved site.

There are a limited number of structurally defined antigenic regions targeted by cross-reactive NA mAbs, and particularly those engaging the CS. Our initial design strategy centered on eliciting CS-directed responses. We therefore used the structurally defined 1G01 and its interaction with NA as a template for glycan placement, as it was the only CS-directed mAb available at the time. However, the more recently defined CS-directed mAbs, namely DA03E17^8^ and FNI9^5^, engage the CS and its periphery with different contacts including distinct angles of approach. Our 1G01- optimized hgNA immunogen, therefore, may have over-restricted access to the CS favoring 1G01- like antibodies while abrogating other antibodies within this CS-directed class. Thus, our observation that there was no demonstrable increase in CS-directed serum responses in mice may be attributed, in part, by CS accessibility. An additional explanation may be due to the relatively long CDR H3 of ∼25 amino acids of the described human CS-directed mAbs^6^. This length is well above the average murine CDR H3 length as well as the average length elicited by the hgNA immunogen, which centered on ∼15 amino acids. Thus, it is likely that a longer CDR H3 is necessary to engage the CS, making the murine model a potential limitation for characterizing CS- focusing immunogens. These observations underscore the importance of having multiple, structurally-defined mAbs that engage a desired epitope (*e.g.,* CS) from different angles and thus different peripheral contacts to guide immunogen design. This will avoid inadvertently occluding conserved epitopes that remain compatible with the intended antibody class, especially when using hyperglycosylation as a design strategy.

Although our initial objective was to enrich CS-directed antibodies, the outcome of our hyperglycosylated immunogen proved informative in an unexpected way. Rather than preferentially enriching CS responses, the hyperglycosylated immunogen elicited antibodies recognizing a previously undefined conserved epitope. This finding suggests that rational glycan engineering may not only reshape immunodominance toward conserved regions but also define antigenic sites that remain underrepresented following conventional vaccine immunization. The interface epitope defined by the Dar1 mAb is reminiscent of the cryptic epitope at the HA head trimer interface recognized by broadly reactive and protective antibodies^25–27,41^. Antibodies engaging this site are non-neutralizing in single-cycle assays but protect in an Fc-dependent manner, suggesting that their mechanism of action involves recognizing HA on an infected cell. Although the HA interface is largely occluded on intact virions, transient conformational dynamics ("breathing") expose this otherwise inaccessible epitope^42^, permitting antibody engagement. While whole virion simulations have not demonstrated HA-like separation of the NA heads^42^, strains of recombinant NA tetramers were shown to adopt “open” conformations that expose the tetramer interface^43^. These observations suggest that transient conformational dynamics permit Dar1 engagement while preserving the structural constraints required for NA tetramer stability. Conservation at the HA interface despite repeated immune exposure has been attributed to such structural constraints^41^, illustrating how cryptic interface epitopes can remain both highly conserved and accessible; it is plausible that the antigenic region recognized by Dar1-like antibodies may be less susceptible to immune escape for similar reasons.

The conserved NA interface has not yet emerged as a defined antigenic site. Since NA is not standardized in seasonal vaccines, repeated vaccine-mediated exposure to this antigenic region is likely limited, leaving natural infection the predominant source of immune exposure. Eliciting Dar1-like antibodies should be a consideration for next-generation influenza vaccines, and the hyperglycosylated NA immunogen described here can serve as a template. The introduced glycans on the hgNA immunogen may shift the conformational equilibrium toward a more open tetramer, more readily exposing the interface epitope even in the presence of a tetramerization domain. While immunogen design approaches have often focused on stabilizing the pre-fusion form of a viral glycoprotein through both covalent and non-covalent modifications, including RSV F^44^, SARS-CoV-2 spike^45^, dengue envelope ^46^, as well as influenza HA^47^ and NA^43^, NA immunogens permitting limited conformational flexibility or full exposure of the tetramer-interface region, such tandemly linked NAs^48^, may more effectively elicit Dar1-like responses.

Several limitations should be considered when interpreting these findings and their implications for NA interface-directed immunity. First, the hgNA immunogen was tested in immunologically naïve mice, which do not reflect the complex influenza immune histories in the human population^49–51^. Second, the prevalence of such NA-interface directed antibodies within the human population, whether following natural infection or vaccination, is not yet known. Thus, it is difficult to assess whether interface-directed NA antibodies in the human population can be or have been elicited, and whether an increase in such antibodies would induce immune pressure against the NA interface. Additionally, although Dar1 mAb conferred partial protection by passive transfer, it is unknown whether direct immunization with the hgNA immunogen would provide comparable protection and whether this translates into broad protection against antigenically diverse influenza viruses. Lastly, because Dar1 did not inhibit NA activity, its protective efficacy is unlikely a result from direct inhibition of CS activity and instead likely depends on Fc-mediated effector functions; further understanding its mechanism of action is necessary. Nevertheless, our NA immunogen provides a framework for leveraging hyperglycosylation to reshape NA immunodominance and enrich antibody responses to conserved antigenic sites.

The variation of seasonal influenza vaccine efficacy and the threat of zoonotic spillover of pre- pandemic influenza viruses motivate the development of next-generation influenza vaccines. Understanding how rationally designed immunogens influence both the serum and single B cell responses can guide iterative design cycles^52,53^. Future design iterations should incorporate the newly-defined tetrameric interface epitope along with other conserved antigenic sites, including the CS and underside. Such epitope-focused immunogen(s) would be a valuable component(s) in next-generation influenza vaccines that aim to elicit broadly protective NA responses.

## MATERIAL AND METHODS

### Study design

The aim of this study was to rationally design NA immunogens using hyperglycosylation to direct humoral responses to conserved sites. We performed *in vivo* murine immunizations and analyzed the serum and single B cell responses. Sample sizes are indicated in the figure legends along with statistical analyses where appropriate.

### NA phylogeny, structural conservation, and glycan modeling

A representative full-length NA strain from each subgroup was submitted to Clustal Omega and analyzed using Multiple Sequence Alignment. The phylogeny tree was generated using FigTree v1.4.4. Structural conservation of the N2 head was computed via ConSurf using 92 H2N2 and 7,576 H3N2 sequence-confirmed isolates from the Bacterial and Viral Bioinformatics Resource Center. Natural and added PNG sites were modeled using high mannose complex hybrid N-glycans on GLYCAM-Web. Historical N2 PNG sites were derived through analysis of PNG sites on 32 H1N2 isolates, 92 H2N2 isolates, 7,334 H3N2 isolates, and 15 H9N2 isolates.

### Expression and purification of recombinant NA antigens

Nucleotide sequences of the wtNA and hgNA versions of the A/Japan/305/1957 (H2N2) tetrameric heads (residues 80-369; N2 numbering) were codon optimized and synthesized by IDT. These gene constructs were cloned into pVRC8400 protein expression vectors with an N-terminal HRV 3C- cleavable 6xHis and Avi-tags, followed by a short “GSG” linker and a non-cleavable hyperglycosylated tetrabrachion domain^39^. Plasmids were sequence-confirmed with Primordium. Monomeric NA head constructs were similarly cloned but without a tetramerization domain. Proteins were transiently expressed in Expi293F cells (ThermoFisher). Five to seven days post-transfection, the supernatants were harvested by centrifugation and purified using cobalt-TALON resin (Takara) affinity chromatography followed by size exclusion chromatography on a Superdex 200 Increase 10/300 GL column (GE Healthcare). Additional NA strains, A/Aichi/2/1968 (H3N2), A/Bilthoven/21438/1971 (H3N2), A/Moscow/10/1999 (H3N2), A/Texas/50/2012 (H3N2), A/Darwin/6/2021 (H3N2), and A/tern/Australia/G70C/1975 (H11N9) were cloned and expressed similarly.

### Expression and purification of IgGs and Fabs

Previously published IgG genes (including the heavy- and light-chain variable domains) were synthesized and codon optimized by IDT, then subcloned into pVRC8400 vector with human constant regions. They were expressed and purified similarly to the NA proteins.

### SDS-PAGE gel electrophoresis

Protein samples were mixed with 10µL of non-reducing Laemmli Sample Buffer (Bio-Rad, Cat#: 1610737). Samples were boiled for 10min at 100°C. 12.5µL of each sample was analyzed on Mini-PROTEAN TGX Stain-Free precast gels (Bio-Rad, Cat#: 456-8026). Gels were then imaged using a ChemiDoc (Bio-Rad). For SDS-PAGE gels under reducing conditions, 2x Laemmli Sample Buffer was mixed with 2-mercaptoethanol prior to mixing with the sample. PNGase F (New England Biolabs, Cat#: P0704S) was added to protein samples according to the manufacturer’s protocol.

### Negative stain electron microscopy

5µl of a given sample (5µg/ml) was adsorbed for 1 minute to a carbon-coated grid (EMS, CF400- CU) that had been made hydrophilic by a 20 second exposure to a glow discharge (25mA). Excess liquid was removed with a filter paper, the grid was floated briefly on water, blotted again, and stained with 0.75% uranyl formate (EMS catalog # 22451). After removing the excess stain with a filter paper, the grids were examined in a TecnaiG² Spirit BioTWIN and images recorded with an AMT NanoSprint43 CCD camera.

### ELISA Assay

Sera and monoclonal antibody reactivity to NA antigens were assayed by ELISA. Briefly, 96-well high binding plates (Corning) were coated with 3-5 µg/ml of NA antigens in PBS at 100µl/well and incubated overnight at 4°C. Plates were blocked with 1% BSA in PBS containing 0.1% Tween- 20 (PBS-T) for 1-2 hours at room temperature (RT). Plates were incubated with serially diluted sera or mAb solutions at RT for 1.5 hours. Sera were serially diluted into PBS with a 10^0.5^ dilution factor (DF), starting from a dilution of 1:40. mAbs were serially diluted by a 10^0.5^ DF into PBS from a starting concentration of 15µg/mL, unless otherwise noted. The final ‘dilution’ for each sample contained PBS alone and was used to calculate background signal for each experiment.

Plates were washed three times with PBS-T. For ELISAs using sera as a primary antibody source, secondary rabbit pAb anti-mouse IgG-HRP (Abcam, AB97046) was added at 1:20,000 dilution in PBS for 1 hour at RT. For mAbs, secondary goat pAb anti-human IgG-HRP (Abcam, AB97225) was added at 1:20,000 dilution in PBS for 1 hour at RT. Plates were washed three times and developed with 1-step ABTS Substrate Solution (ThermoFisher) for 45 minutes, then stopped with 100uL of 1% SDS solution. Absorbance was measured using a plate reader at 405nm. EC_50_ values were determined by non-linear regression (sigmoidal) using GraphPad Prism 10.2.3 software. Endpoint sera titers were defined as the dilution value just above an OD_405nm_ cutoff.

### ELLA Assay

96 well plates were coated with 25µg/mL of fetuin (Sigma Aldrich, F3004) and kept at 4°C overnight. Plates were blocked with PBS, 1% BSA, 0.1% Tween-20 solutions and put on a shaker for 2 hours at RT. Plates were then washed 6x with ELISA buffer. Recombinant NA_TB_ was serially diluted twofold in DPBS containing Ca²⁺/Mg²⁺ (Gibco, Cat#:14040117), beginning at 0.005 µg/mL, and 100 µL of each dilution was added to the plate; DPBS alone served as the negative control. Plates were incubated at 37°C for 2 hours and then washed. Plates were incubated with 100µL of either 1 or 5µg/mL of HRP-conjugated lectin from Arachis hypogaea (peanut) (Sigma Aldrich, L7759) for 75-100min at RT on a plate shaker. Plates were then washed and coated with 100µL of ABTS and imaged imaged. EC_50_ values of NA tetramers were determined by non-linear regression (sigmoidal) using GraphPad Prism 10.2.3 software.

### MUNANA Assay

Small substrate MUNANA fluorescence assays were performed as described^54^. Recombinant NA tetramer was serially diluted in using 1:2 dilutions from a starting concentration of 25µg/mL with a buffer-only control included. 50µL of the serially diluted NA were transferred to black 96-well flat-bottom plates. 50µL of 300µM MUNANA solution was added per well and the plate was incubated at 37°C for 1 hour. 300µM of MUNANA substrate was prepared using 2’-(4- methylumbelliferyl)-a-D-N-acetylneuraminic acid (Sigma Aldrich, M8639). Fresh substrate was prepared for each experiment. After 1 hour, 100µL of stop solution using NaOH was added per well, and read using 355nm excitation and 460nm emission wavelengths.

### MUNANA inhibition assay

wtNA_TB_ protein was prepared at 3xEC_50_ in assay buffer, and 50µL of the solution was distributed into each well of a black 96-well. Sera from each mouse were diluted to 3x (1/40) in a separate plate. 50µL of the 3x mixture of serum dilution was added to the 96-well plate and incubated at RT for 1hr. 50µL of 300µM MUNANA assay were added to each well, and the plate incubated at 37°C for 1 hour. 100µL of stop solution was then added, and plates were read using 355nm excitation and 460nm emission wavelengths. For mAb MUNANA inhibition, mAbs were serially diluted instead of sera. Data in the figures reflect effective concentrations.

### ELLA inhibition assay

96 well plates were coated with fetuin overnight at 4°C and blocked as described for ELLA assays^55^. For the primary incubation, 65µL of 2x the EC_50_ enzyme activity concentration (for a given recombinant NA tetramer) was premixed with 65µL of 2x serial dilution of either a mAb or mouse sera sample. Plates were incubated at 37°C incubator for 2hrs and followed the ELLA assay protocol described above. IC_50_ values of a given mAb or sera were calculated using a non-linear regression model.

### Immunizations

Immunizations were performed using female C57BL/6 mice purchased from The Jackson Laboratory (Bar Harbor, ME) aged 8-10 weeks. Mice received 100µL of inoculum intraperitoneally, containing 20µg of protein adjuvanted with 50% v/v Sigma adjuvant in sterile PBS. Mice were bled from the submandibular vein on day -1 and received their first immunization on day 0. Boosts occurred on day 21 and day 42. For the wtNA_TB_ and hgNA_TB_ experiment, mice were bled at day 21 and day 42 prior to receiving boosts. Experiments were conducted with institutional IACUC approval (2014N000252).

### Competition ELISA

Sera competition ELISAs with recombinant monoclonal antibodies were performed with hgNA_TB_ or Dar’21 NA antigens. Briefly, 3µg/mL of antigen was coated onto the plate and incubated overnight at 4°C. After blocking, plates were incubated with a 10^0.5^ serial dilution of mAbs at a starting concentration of 50µg/mL (except for hg23 at 15µg/mL) and incubated at RT for 1hr. 10µL of sera solution from hgNA_TB_-immunized mice was then added to each well such that the final concentration of sera was at the EC_50_ of the serum ELISA titration curve for the respective coating antigen. The primary mAbs were at an effective concentration of 40 (or 12) µg/mL at their highest concentrations. Sera and mAb were incubated on the antigen-coated plates for 1 hour. Plates were then washed 3x in PBS-T and incubated for 1hr with HRP-conjugated goat anti-mouse IgG, human/bovine/horse SP ads antibody (Southern Biotech) at a 1:8000 dilution. Plates were washed 3x with PBS-T and developed with 1-step ABTS for 45 minutes, then stopped with 1% SDS solution and measured for 405nm. Competition EC_50_ values were determined by non-linear regression (sigmoidal) using GraphPad Prism 10.2.3 software. Percent competition was determined by calculating [1-(OD_405nm_ at max [mAb] wells / OD_405nm_ of PBS-only wells) x 100] using GraphPad Prism 10.2.3 software. Sera from day 0 was used as a negative control and percent competition was calculated relative to the no-IgG control wells.

### Competition biolayer interferometry (BLI)

NDS.1, NDS.3, and Dar 1 Fabs and Dar’21 NA monomer were recombinantly expressed as described above. For competition experiments, the C-terminal purification tags were cleaved from the NA and Fab constructs and were pre-incubated at a 1:1.2 molar stoichiometry, respectively. Pre-bound NA:Fab complexes were then added as the analyte to tagged Fab (0.2mg/ml) bound to Ni-NTA biosensor. Self-competition with each Fab was used as reference relative to response obtained from Dar’21 NA with no pre-bound Fab binding to tagged Fab on the biosensor; successful competition is reflected as a loss of response (nm).

### Probe generation

Avi-tagged monomeric and tetrameric wtNA head constructs were biotinylated using a BirA biotin-protein ligase kit (Avidity) according to manufacturer’s protocol. The biotinylated proteins were subsequently repurified by size exclusion chromatography on a Superdex 200 Increase 10/300 GL column (GE Healthcare) in PBS.

### Single B cell sorting

Spleens from wtNA-immunized and hgNA-immunized mice were harvested on d56 for single B cell sorting. A single cell suspension was generated by passing spleen tissue through a 70µm strainer and washed with FACS buffer (PBS, 25mM HEPES, 1% heat-inactivated FBS). The single cell suspension was then centrifuged for 10 mins at 4°C and 1300 RPM. Red blood cells were lysed using 2mL ACK lysis buffer for 2-3mins on ice, then quenched with FACS Buffer. Clarified cell pellets were washed 2x with FACS buffer. For the sort described in **Fig. 4**, two samples from each cohort were stained for single B cell sorting. The remaining 3 samples from each cohort were frozen in HI-FBS with 10% DMSO for later use. Cells were stained for the following cell markers for flow cytometry: CD3e – AF700, CD19 – BV605, msIgG -BV421, msIgM - PerCP/Cy5.5 at a 1:100 dilution. Biotinylated antigens used to stain cells *from wtNA-immunized mice* were labeled with streptavidin-conjugated fluorophores as follows: wtNA_TB_ – PE, wtNA_TB_ – APC, hgNA_TB_ – APC/Cy7. Individual Live/CD19+/IgG+/ wtNA_TB_-PE+/ wtNA_TB_-APC+ cells were sorted and collected into 96-well plates with 4µL of lysis buffer (0.5xPBS, 10 mM DTT, and 4 units of RNaseOUT (ThermoFisher)). Biotinylated antigens used to stain cells from *hgNA_TB_-immunized mice* were labeled with streptavidin-conjugated fluorophores as follows: hgNA_TB_ – PE, hgNA_TB_ – APC, wtNA_TB_ – APC/Cy7. These fluorophores were added at a final concentration of 25nM in FACS buffer and incubated with splenocytes for 45mins-1hr on ice. Cells were then washed with cold FACS buffer and stained with Live/Dead aqua in cold PBS for 15-20 mins. Cells were washed and brought to a volume of 1.5mL in FACS buffer for sorting. Individual Live/CD19+/IgG+/ hgNA_TB_-PE+/ hgNA_TB_-APC+ cells were sorted and collected into 96-well plates with 4uL of lysis buffer. For the sort represented with **Fig. 5**, frozen splenocytes from two mice in each cohort were thawed slowly in 10mL of pre-warmed RPMI media with 50% HI-FBS. The cells were washed with FACS buffer and stained with the following cell markers for flow cytometry: CD3e – AF700, CD19 – BV605, msIgG – BV421, msIgM – PerCP/Cy5.5 at a 1:100 dilution. Biotinylated antigens used to stain cells from all mice samples were labeled with streptavidin-conjugated fluorophores as follows: Dar’21 - PE, Dar’21 - APC, wtNA_TB_ - FITC, wtNA_TB_ - APC/Cy7, hgNA_TB_ - PE/Cy7, hgNA_TB_ - BV605. These fluorophores were also added at a final concentration of 25nM in FACS buffer and incubated with splenocytes for 45mins-1hr on ice. Cells were then washed with FACS buffer stained with Live/Dead aqua in PBS. Individual Live/CD19+/IgG+/Dar’21-PE+/Dar’21- APC+ cells were sorted and collected into 96-well plates with 4µL of lysis buffer. All biotinylated NAs were individually premixed with fluorescently labeled streptavidin (SA) at a 4:1 molar ratio for 30min at 4°C prior to sorting to form fluorescent antigen tetramers. 3-6 million events were recorded for each sample for antigen-specific frequency determination, and the remainder of the sample was sorted as single cells. Plates were centrifuged at 3000 x g for 1 minute and stored at - 80°C before reverse transcription and V(D)J sequencing. Flow cytometry data was analyzed using FlowJo software, version 10.10.0.

### B cell receptor sequencing

Thawed B cell lysates were reverse transcribed using the SuperScriptIV VILO MasterMix (ThermoFisher) in a total volume of 20µL according to the manufacturer’s recommendations. Two rounds of PCR, performed separately for the heavy and light chains, were then executed using previously published primers^56,57^ and the Herculase II Fusion Enzyme, according to manufacturer’s recommendations (Agilent, Cat# 600679). Variable heavy and light chains were analyzed by agarose gel electrophoresis, sequenced via Sanger sequencing (Azenta), and analyzed using IMGT High V-Quest. For reads with low-quality terminal sequence, up to 30 ambiguous nucleotides at the 5′ or 3′ end were replaced with the corresponding inferred germline sequence for cloning purposes. For genetic analysis, only high quality ‘productive’ reads with defined CDRs were included. Paired heavy- and light-chain variable-region sequences from cells yielding high-quality reads were codon-optimized for mammalian cells (IDT).

### Cryo-EM sample preparation and data collection

The H2N2 Japan/1957 NA head monomer was incubated with the 1G01 and Dar1 Fab at a molar ratio 1:2:2. The monomer-Fab complexes were then purified using size exclusion chromatography on a Superdex 200 Increase 10/300 GL column (GE Healthcare). Fractions with stoichiometric complexes were concentrated to 0.5-1 mg/mL for application onto cryoEM grids. Quantifoil R 1.2/1.3 (Cu, 300-mesh; Quantifoil Micro Tools GmbH) grids were used and treated with Ar/O_2_ plasma (Solarus plasma cleaner, Gatan) for 8s before sample application. Grids were prepared using a Vitrobot Mark IV (Thermo Fisher). n-Dodecyl-β-D-Maltoside (DDM; Anatrace) at final concentration of 0.06 mM was used. Grids were plunge-frozen into liquid ethane.

### Cryo-EM data collection

Grids were imaged using a Titan Krios microscope, operated at 300 kV and equipped with a Gatan K3 Summit direct electron detector. A total of 27,430 movies were collected for a total dose of 52.5 e−/Å^2^. Images were collected at a nominal magnification of 165K, corresponding to a calibrated pixel size of 0.736 Å/pixel. EPU software (Thermo Fisher) was used for automated data collection. The image processing for the Dar1 complex is shown in **fig. S10**. In brief, dose fractionated images were motion corrected with the RELION^58^ implementation of MotionCor followed by CTF estimation in cryoSPARC Patch CTF. Particle picking was carried out using template matching in cryoSPARC using a 20Å low pass filter, resulting in 4,364,717 particles. Initial steps were carried out with particles downsampled to ∼1.56 Å/pixel. Particles underwent one round of 2D classification to remove false positives from particle picking, resulting in 2,166,833 particles. 3D classification yielded 838,082 particles which were extracted at full resolution and underwent particle polishing in RELION to produce a ∼2.8 Å reconstruction after blush refinement in RELION (**table S1**).

### Model building and refinement

The initial Dar1:1G01:N2 NA model was obtained by docking a Dar1 Fab (generated using AlphaFold^59^) and N2 NA JP’57 (from PDB 3TIA) and 1G01 Fab (from PDB 6Q23) in ChimeraX^60^. Multiple rounds of real-space refinement in Phenix^61^ using reference model restraints followed by manual adjustment in Coot^62^; the refined model was validated with MolProbity^63^ (**table S1**). Structural analysis and figures were done using PyMol. Structural biology applications used in this project were compiled and configured by SBGrid^64^.

### Passive transfer of monoclonal NA antibodies and viral challenge

mAbs were infused at 5 mg/kg intraperitoneally in 8-10 weeks old wildtype C57BL/6 mice (n=10 mice per group, including 5 male and 5 female (Jackson Laboratory). Two hours later, the mice were intranasally infected with 100% lethal dose of H3N2 X-31 (BEI Resources cat# NR-3483) (10^8^ TCID_50_/ml)^65^. The mice were monitored for 21 days for body weight loss and survival. H3N2 X-31 viruses were cultured in MDCK cells and quantified by TCID_50_ in MDCK cells^66^. All experiments were conducted with institutional IACUC approval (2014N000252).

### Statistics

Statistical analyses used are noted in the corresponding figure legend.

## SUPPLEMENTARY MATERIALS

figures S1-S11 table S1

## Acknowledgements

We thank Maria Ericsson and the Harvard Electron Microscopy Core for collection of negative stain images. We thank Richard Walsh and the Harvard Cryo-EM Center staff for their assistance with cryo-EM data collection. We thank Jared Feldman, Emerson Glassey, Isaiah Shriner, Anne Roffler, and Goran Bajic for helpful discussions as well as Steve Harrison for critical reading of the initial manuscript.

## Funding

We acknowledge support from the following sources. P01 AI089618 (AGS).

F30 AI181355 (RH).

R01AI153098, R01AI155447, R01AI195539 (DL).

We acknowledge support from NIGMS T32 GM0008313.

This research has been funded in whole or part with federal funds under a contract from the National Institute of Allergy and Infectious Diseases, NIH contract 75N93019C00050 (AGS).

The content is solely the responsibility of the authors and does not necessarily represent the official views of the National Institutes of Health.

We acknowledge the Harvard University Biophysics Graduate Program.

## Author contributions

Conceptualization: RH, AGS

Methodology: RH, FNMA, SZ, SR, LR, DTL, TC

Investigation: RH, FNMA, DTL, SEL, CLM, SR, SZ, LR,

Visualization: RH, FNMA, SZ, SR, AGS

Funding acquisition: RH, DL, AGS

Project administration: DL, AGS

Supervision: DL, AGS

Writing – original draft: RH, AGS

Writing – review & editing: RH, FNMA, SZ, SR, SEL, CLM, DTL, LR, TC, DL, AGS

## Data and materials availability

Data to support the conclusions of the paper can be found in the Supplementary Materials. Antibody sequences are available at GenBank under accession codes PZ760929-PZ761094 (wt and hgNA-elicited) and PZ761095-PZ761152 (Dar’21 NA-enriched). The map of the cryo-EM reconstruction of 1G01 and Dar1 Fabs bound to NA is available at the Electron Microscopy Data Bank (accession number EMD-78520). Coordinates for the refined model of 1G01:Dar1:NA complex is available at the Protein Data Bank (accession number 37VO). Requests for material should be addressed to Aaron G. Schmidt.

