## Supplemental Information for "A rationally designed neuraminidase immunogen elicits humoral responses to a conserved viral site"

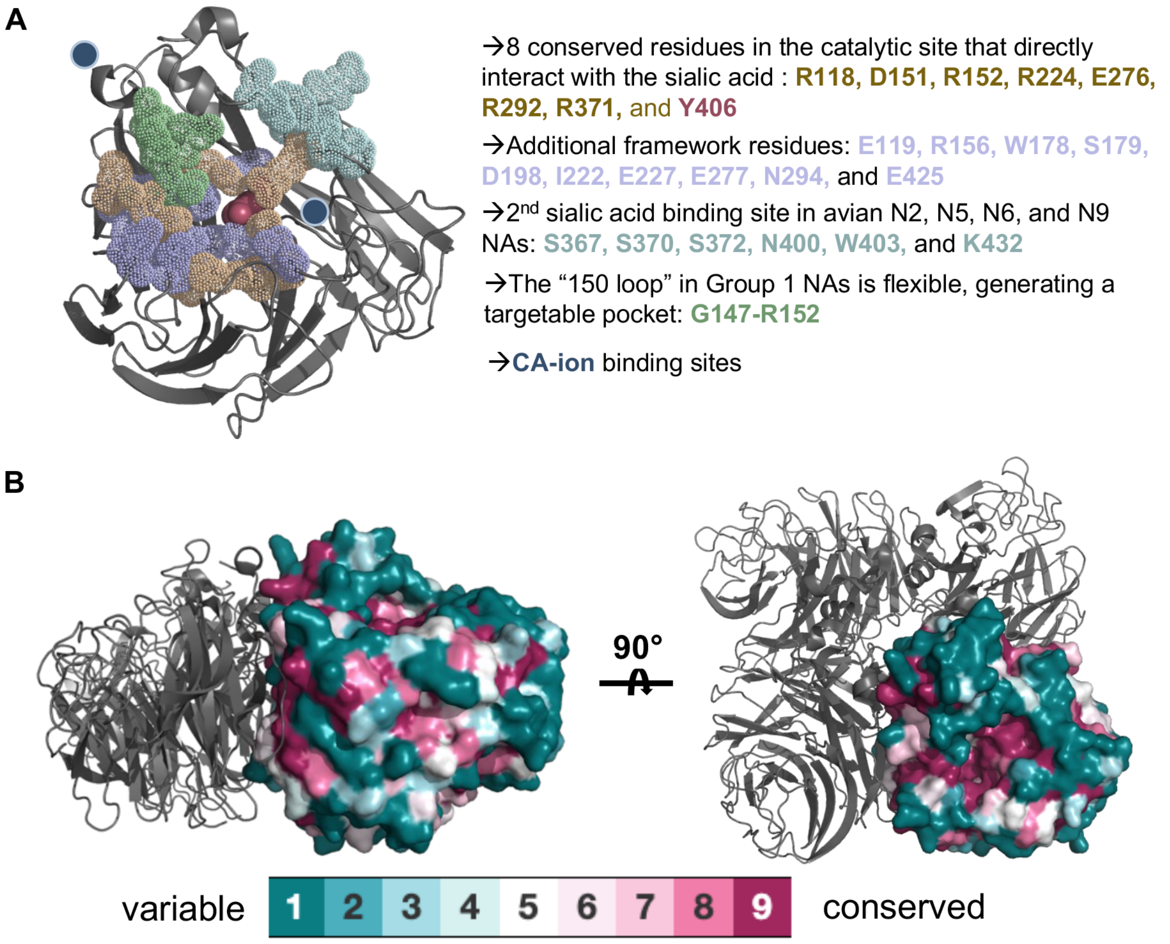

**fig. S1: Neuraminidase structural conservation. (A)** A/Japan/305/1957 (J'57) H2N2 monomer (PDB 6Q20) shown in gray cartoon. Conserved amino acid residues are shown as color-coded spheres. Each monomer has at least one calcium ion site and there is a calcium ion site in the center of the tetramer, shared between the protomers. **(B)** N2 structural conservation shown on the J'57 N2 NA tetramer (PDB 6Q20), produced using ConSurf by analyzing 92 H2N2 and 7,576 H3N2 sequence-confirmed isolates from the Bacterial and Viral Bioinformatics Resource Center.

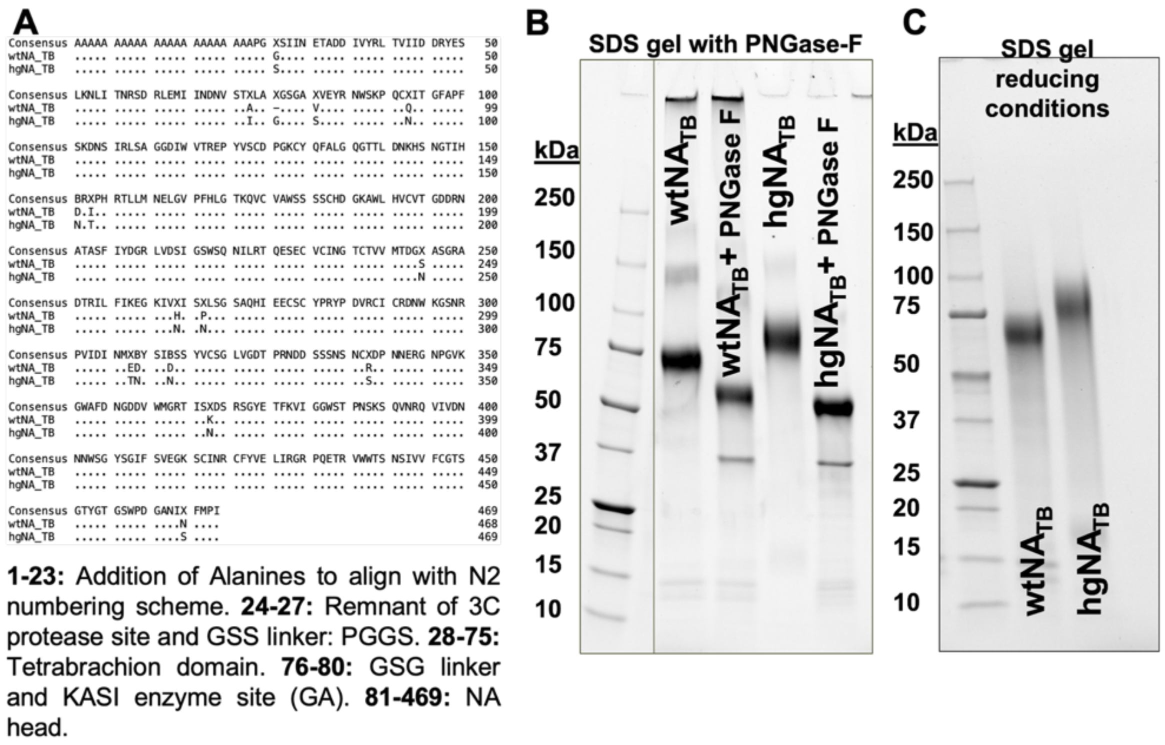

**fig. S2: Characterization of wtNA$_{TB}$ and hgNA$_{TB}$ proteins. (A)** Amino acid sequence alignment between wtNA and hgNA constructs using N2 numbering from A/Japan/305/1957. 10 historical and one novel predicted N-linked glycosylation (PNG) sites were added to make the hgNA. **(B)** SDS-PAGE analysis of PNGase treated proteins and an equivalent amount of non-treated protein. **(C)** SDS-PAGE analysis of proteins under reducing conditions.

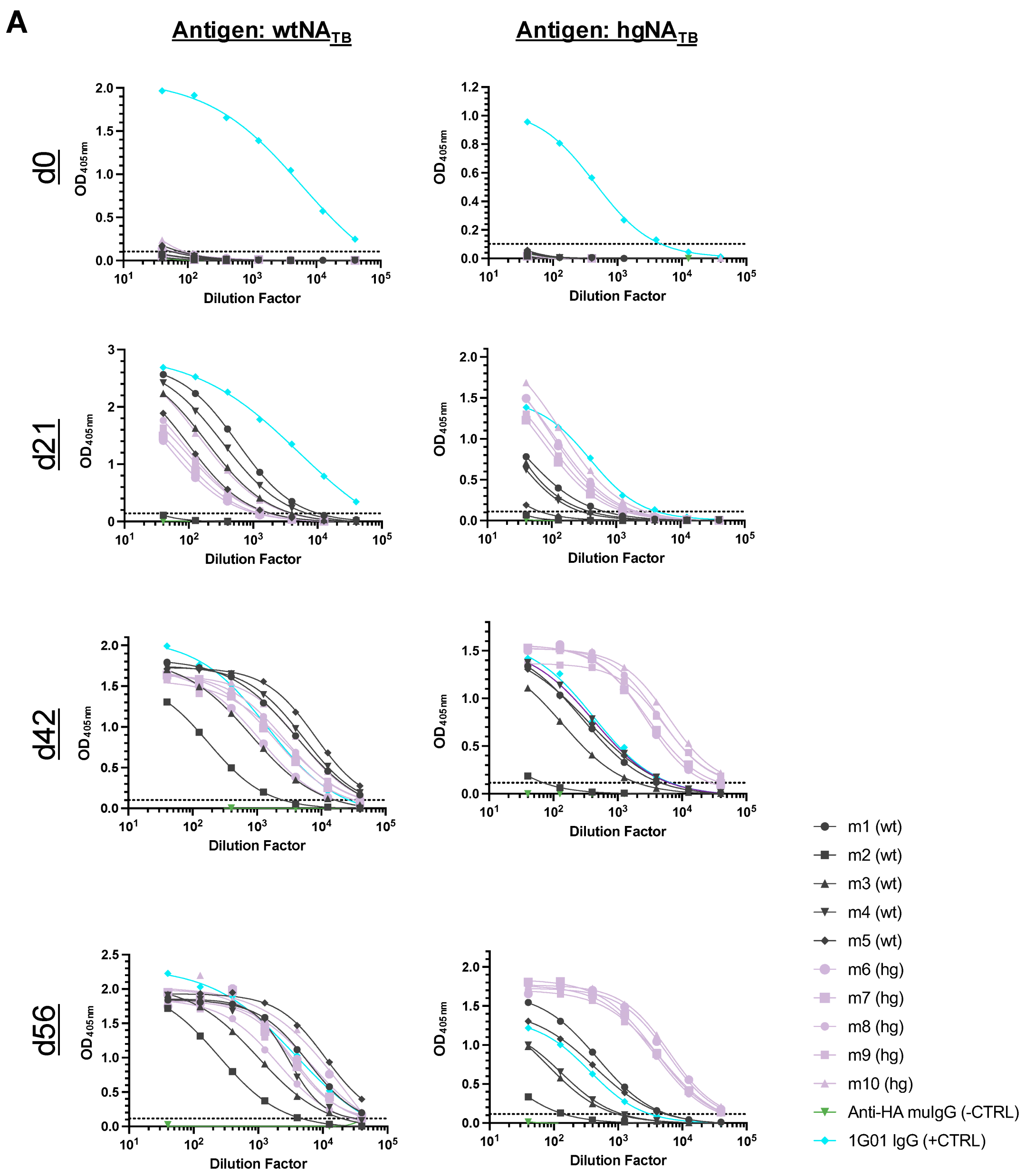

A
Antigen: wtNA$_{TB}$
Antigen: hgNA$_{TB}$
d0
d21
d42
d56
OD$_{405nm}$
Dilution Factor
m1 (wt)
m2 (wt)
m3 (wt)
m4 (wt)
m5 (wt)
m6 (hg)
m7 (hg)
m8 (hg)
m9 (hg)
m10 (hg)
Anti-HA muIgG (-CTRL)
1G01 IgG (+CTRL)

**(continued)**

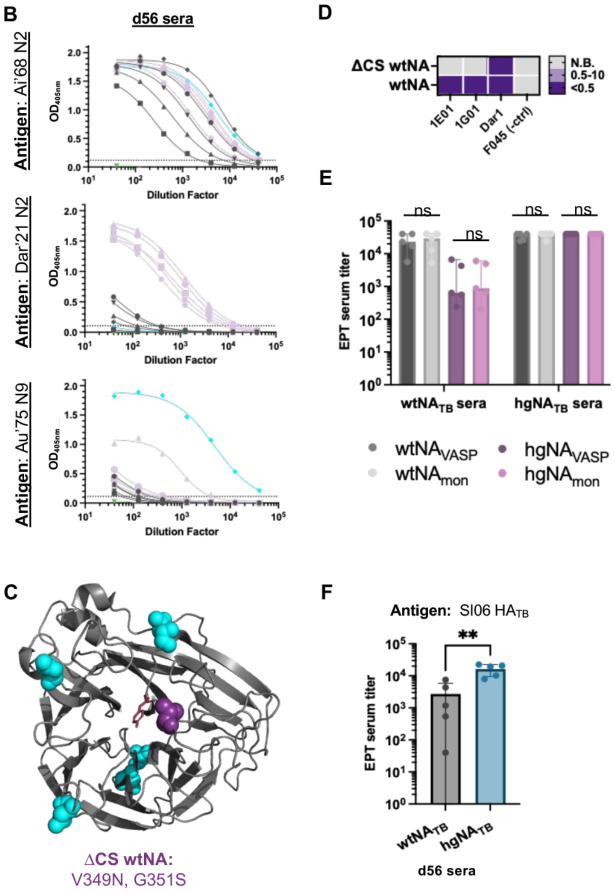

**fig. S3. Sera-focusing in the hgNA$_{TB}$ immunized cohort. (A)** Sera ELISA reactivity of d0, 21, 42, and 56 to NA proteins. Sera from mice immunized with hgNA$_{TB}$ (pink) or wtNA$_{TB}$ (gray) immunogens. The dotted line in each graph represents the threshold for the endpoint serum dilution factor (EPT DF), which was calculated by taking the average OD$_{405nm}$ signal from all blank wells and adding 2x the standard deviation of the OD$_{405nm}$ signal from all blank wells. **(B)** d56 sera reactivity from wtNA$_{TB}$-immunized mice (gray) and hgNA$_{TB}$-immunized mice (pink) against historical NA monomers. **(C)** The ΔCS wtNA included an engineered PNG site in the CS pocket using V349N and G351S mutations. V349 is shown in purple spheres. Cyan spheres show wildtype PNG sites. **(D)** Reactivity of mAbs against the ΔCS wtNA in ELISA. **(E)** Reactivity to monomeric versus VASP-tetramerized wtNA and hgNA proteins showed no differences. The VASP tetramerization tag was used to isolate differences in reactivity to the NA-head specifically as opposed to possible differences due to TB-directed responses. Data was compared using a Mann-Whitney test. **(F)** Sera reactivity of wt- and hgNA$_{TB}$-immunized mice to the TB tetramerization domain with an H1 Solomon Islands/03/2006 HA head. Data was compared using a Mann-Whitney test, p-value: **0.0079. Bars represent mean with standard deviation.

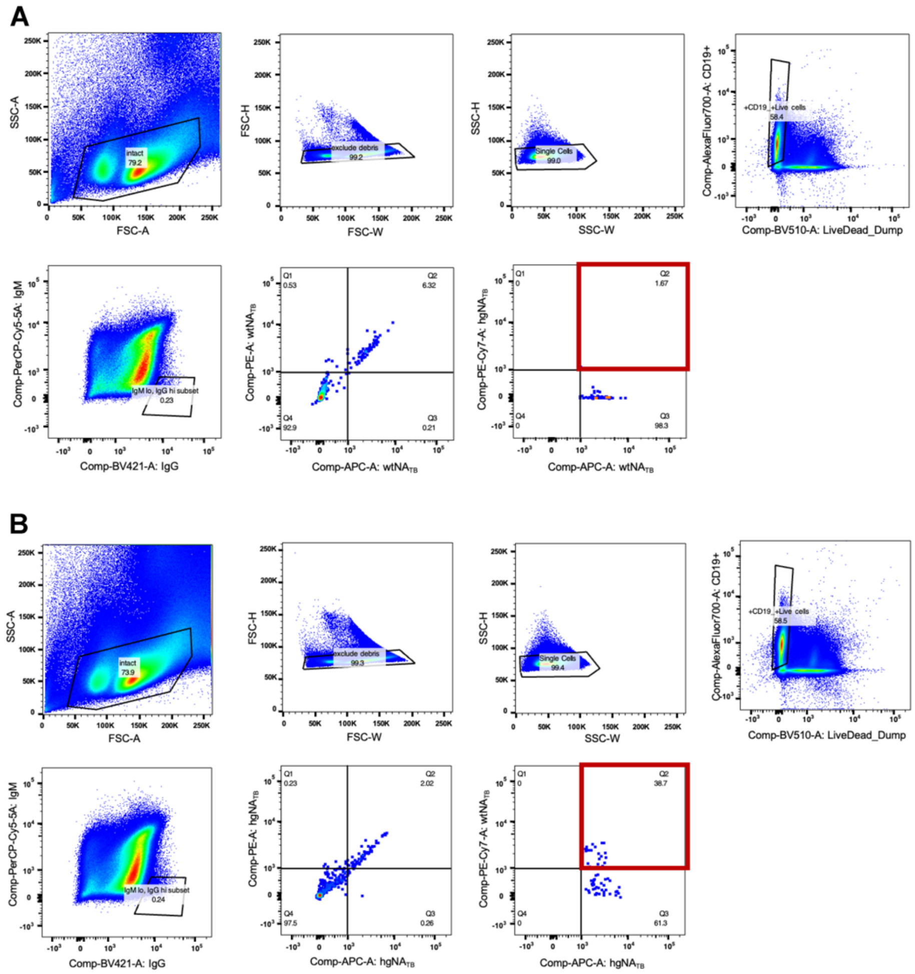

**fig. S4. Flow cytometry gating strategy for wt- and hgNA- immunized mice. (A)** Representative sample from wtNA$_{TB}$-immunized mouse (n=2 wtNA$_{TB}$-immunized spleens, each divided into two cell samples). Splenocytes from wtNA$_{TB}$ B cells were gated on CD19+, IgM$^{low}$, and IgG$^{high}$, and wtNA$_{TB}$ antigen specificity before being index sorted. Red box indicates the lack of orthogonal reactivity for hgNA$_{TB}$ in the wtNA$_{TB}$-specific positive cells. **(B)** Representative sample from hgNA$_{TB}$-immunized mouse (n=2 hgNA$_{TB}$-immunized spleens, each divided into two cell samples). An identical gating strategy to wt was used for splenocytes isolated from hgNA$_{TB}$-immunized mice. Antigen-specific cells showed substantially greater recognition of the orthogonal wtNA immunogen (red box). hgNA$_{TB}$ antigen specific cells were index sorted for further analyses.

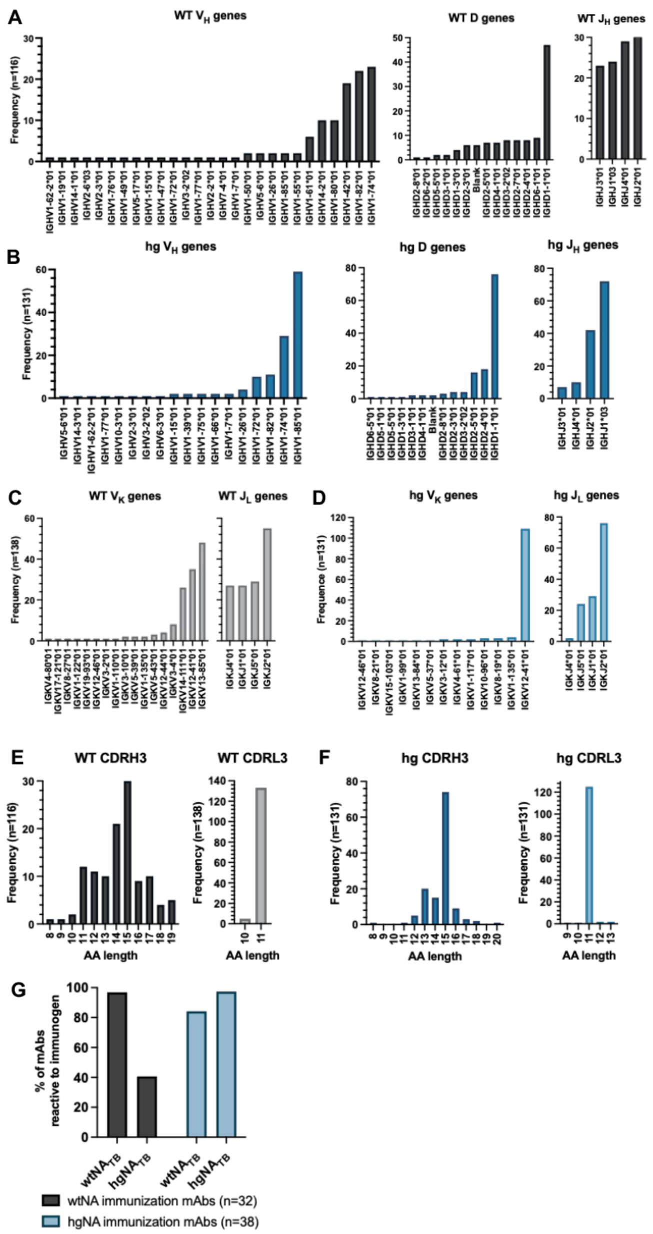

**fig. S5. Genetic analyses and features of splenic B cells from wtNA$_{TB}$- and hgNA$_{TB}$- immunized mice.** **(A)** V-, D-, and J- heavy chain genes from wtNA$_{TB}$-immunized mice. **(B)** V-, D-, and J- heavy chain genes from hgNA$_{TB}$-immunized mice. **(C)** V- kappa and J- light chains from wtNA$_{TB}$-immunized mice. **(D)** V- kappa and J- light chain genes from hgNA$_{TB}$-immunized mice. CDR H3 and CDR L3 amino acid lengths of sequenced B cells from **(E)** wtNA$_{TB}$- and **(F)** hgNA$_{TB}$-immunized mice. **(G)** Reactivity profiles of mAbs derived from wtNA$_{TB}$- and hgNA$_{TB}$-immunized mice. Statistical significance was evaluated using Mann-Whitney tests, with p-value: ****<0.0001. All alleles are *01, unless indicated otherwise.

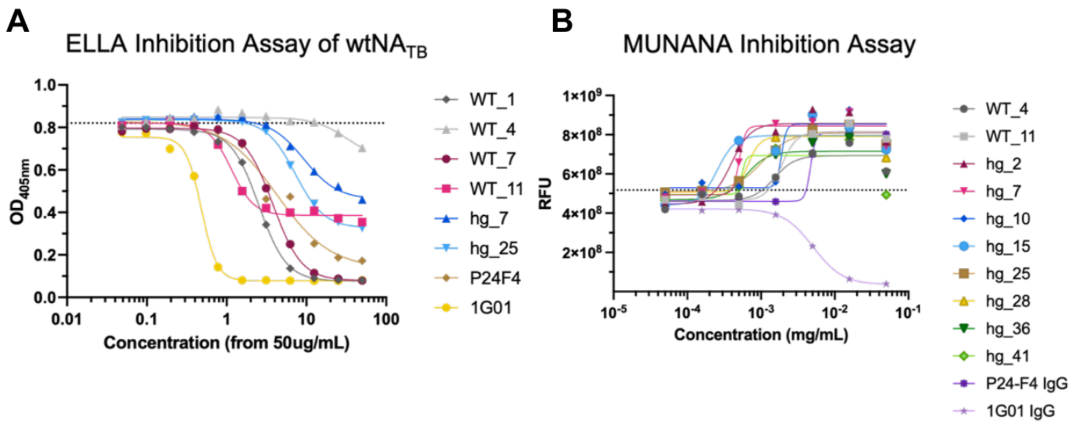

**fig. S6. Inhibition of NA enzymatic activity by isolated mAbs. (A)** ELLA and **(B)** MUNANA inhibition assays using mAbs derived from isolated single B cells. 1G01 and P24F4 (an in-house N2-reactive mAb) IgGs are used as positive controls.

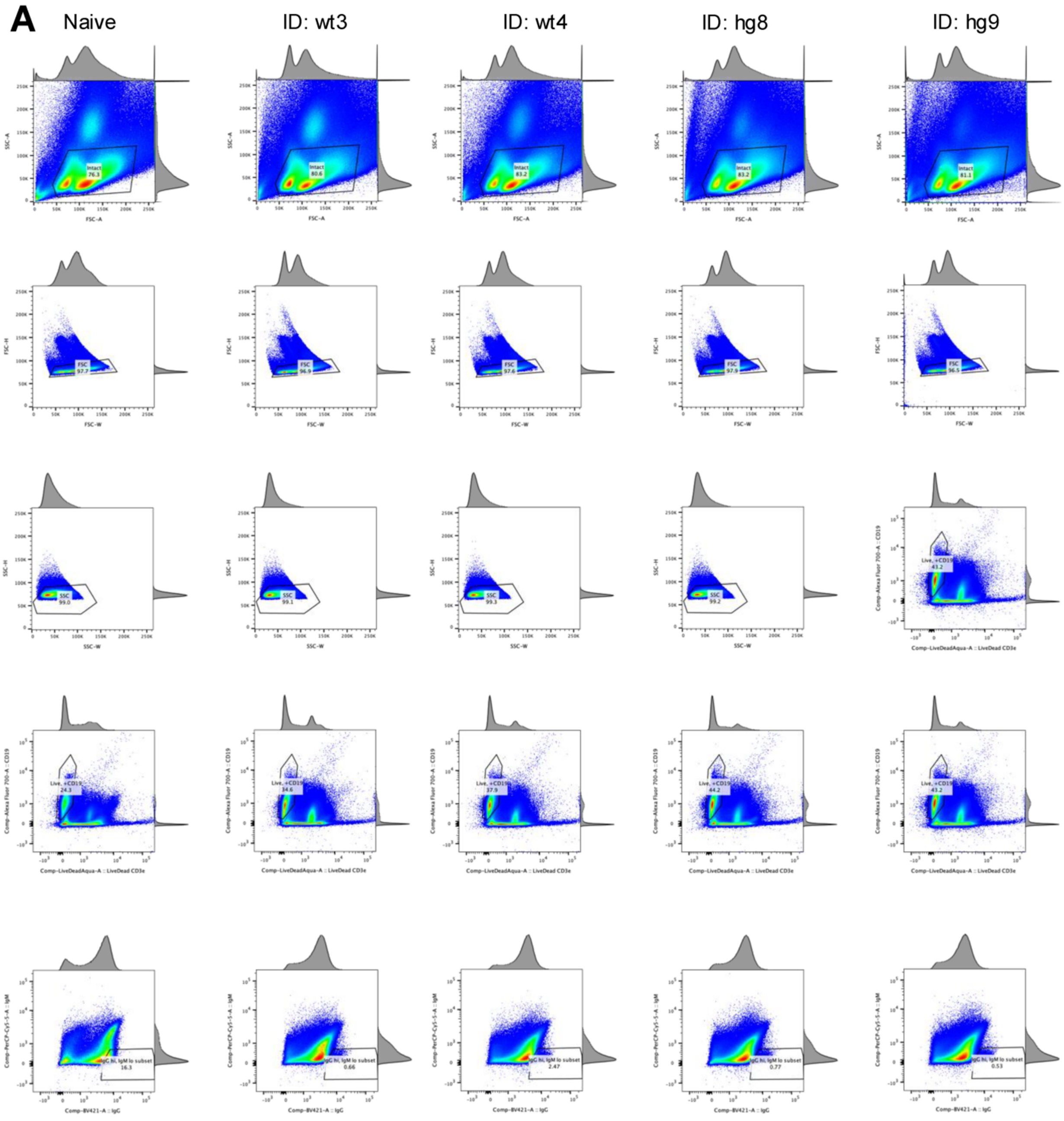

A
Naive
ID: wt3
ID: wt4
ID: hg8
ID: hg9
Intact 76.3
Intact 80.6
Intact 83.2
Intact 83.2
Intact 81.1
SSC-A
FSC-A
FSC 97.7
FSC 96.9
FSC 97.6
FSC 97.9
FSC 96.5
FSC-H
FSC-W
SSC 99.0
SSC 99.1
SSC 99.3
SSC 99.2
SSC-H
SSC-W
Live, +CD19 43.2
Live, +CD19 24.3
Live, +CD19 34.6
Live, +CD19 37.9
Live, +CD19 44.2
Live, +CD19 43.2
Comp-Alexa Fluor 700-A :: CD19
Comp-LiveDeadAqua-A :: LiveDead CD3e
IgG hi, IgM lo subset 16.3
IgG hi, IgM lo subset 0.66
IgG hi, IgM lo subset 2.47
IgG hi, IgM lo subset 0.77
IgG hi, IgM lo subset 0.53
Comp-PerCP-Cy5-5-A :: IgM
Comp-BV421-A :: IgG

**(continued)**

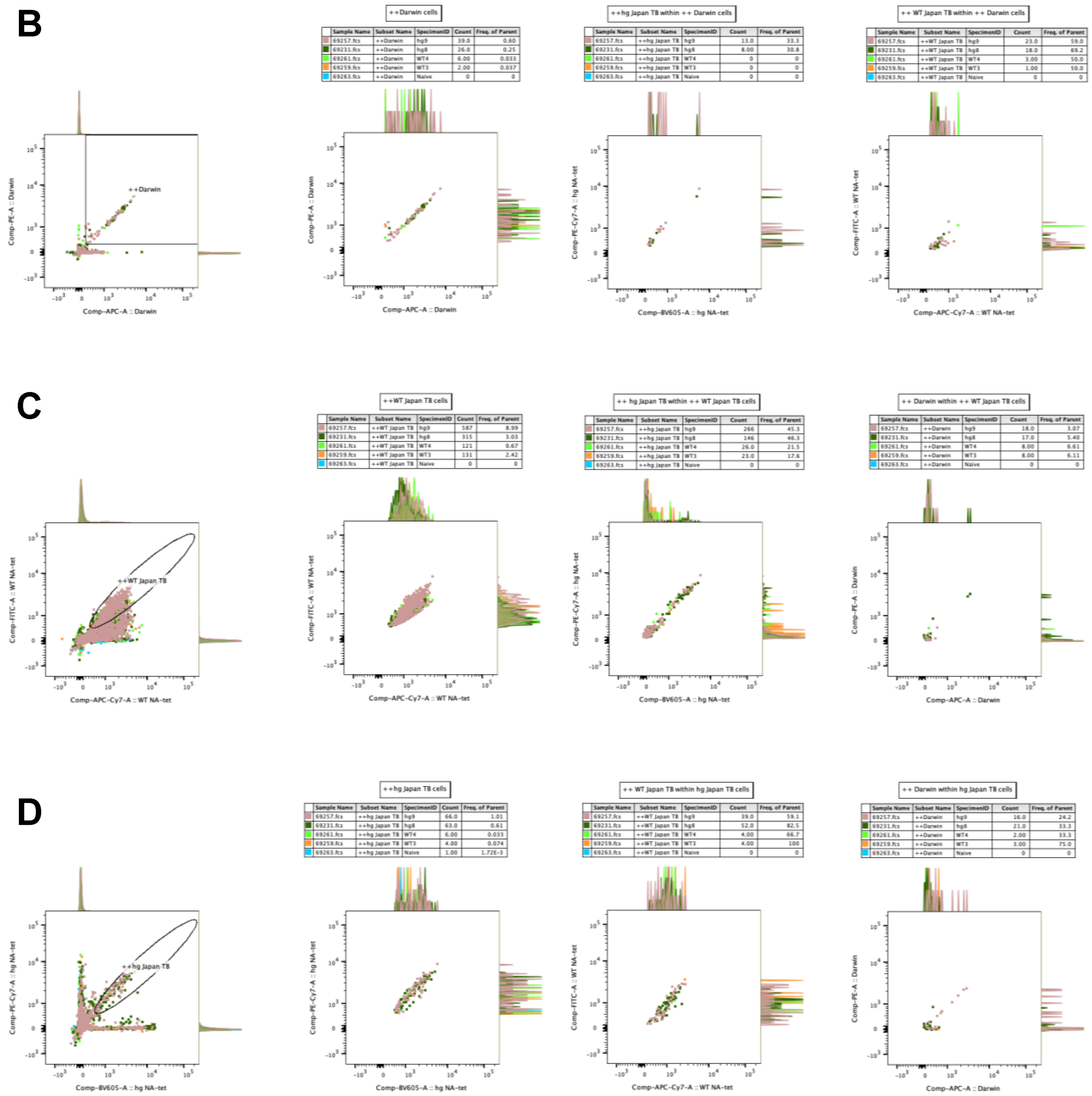

**fig. S7. Flow cytometry data from sort using Dar'21 N2 NA as a sorting probe. (A)** Splenocytes from: naïve mouse (n=1), wtNA$_{TB}$-immunized mice (n=2), and hgNA$_{TB}$-immunized mice (n=2) were gated on live, CD19+, IgM$^{low}$, IgG$^{high}$ before being analyzed for reactivity to **(B)** Dar'21, **(C)** wtNA$_{TB}$, and **(D)** hgNA$_{TB}$ proteins. Each antigen was conjugated to two unique fluorophores. B cells that were double-positive for Dar'21 NA were index sorted.

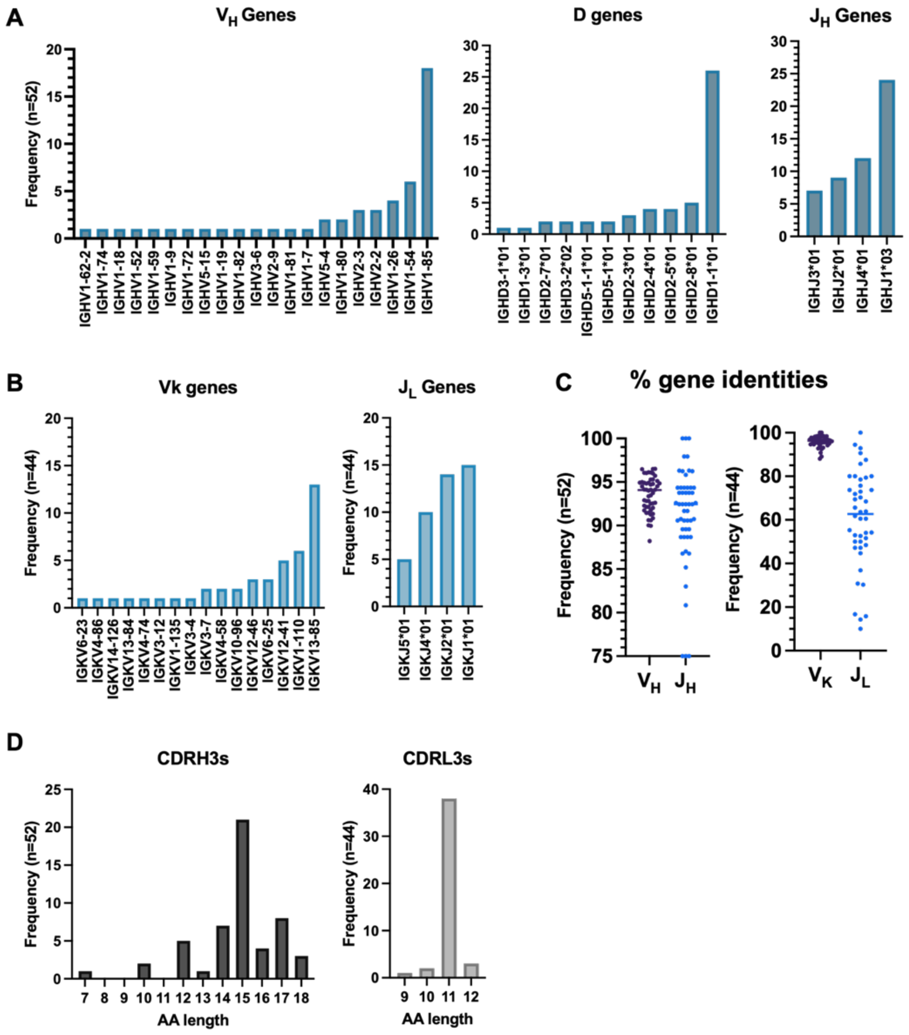

**fig. S8. Genetic analyses of splenic B cells isolated from $hgNA_{TB}$-immunized mice using Dar'21 NA.** **(A)** V-, D-, and J- heavy chain gene analysis. **(B)** V- kappa and J- light chain analysis. All alleles are *01 unless indicated otherwise. **(C)** Percent sequence identities of B cells to germline. The mean identity percents of $V_H$, $J_H$, $V_K$, and $J_L$ were ~93.5%, ~91.2%, ~95.9%, and ~60.5%, respectively. **(D)** CDR H3 and CDR L3 length analysis.

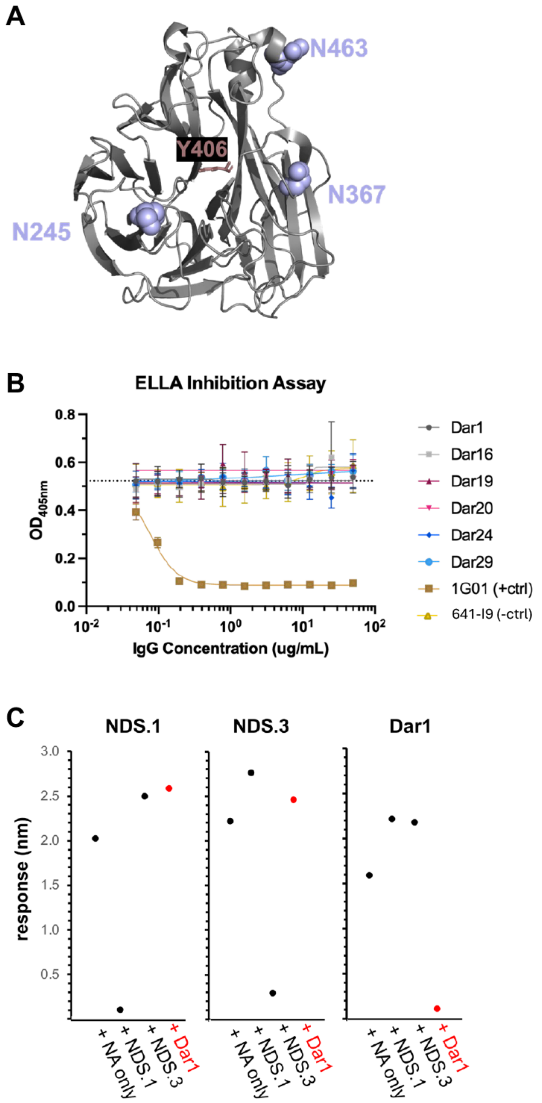

**fig. S9. Dar'21 NA glycan variants and additional characterization of Dar'21-reactive mAbs. (A)** Surface locations of 3 PNGs that differ between Dar'21 and wtNA NA strains. Each glycan site was individually mutated to the wtNA amino acids and tested with Dar1 mAbs to determine glycan dependencies. **(B)** ELLA inhibition assay of a subpanel of mAbs from the Dar'21-reactive sort. **(C)** Competition-based BLI using non-overlapping underside NDS.1 and NDS.3 Fabs for crude epitope mapping. Each panel shows binding to Dar'21 NA only and competition with respective Fabs. Data are representative of two independent experiments.

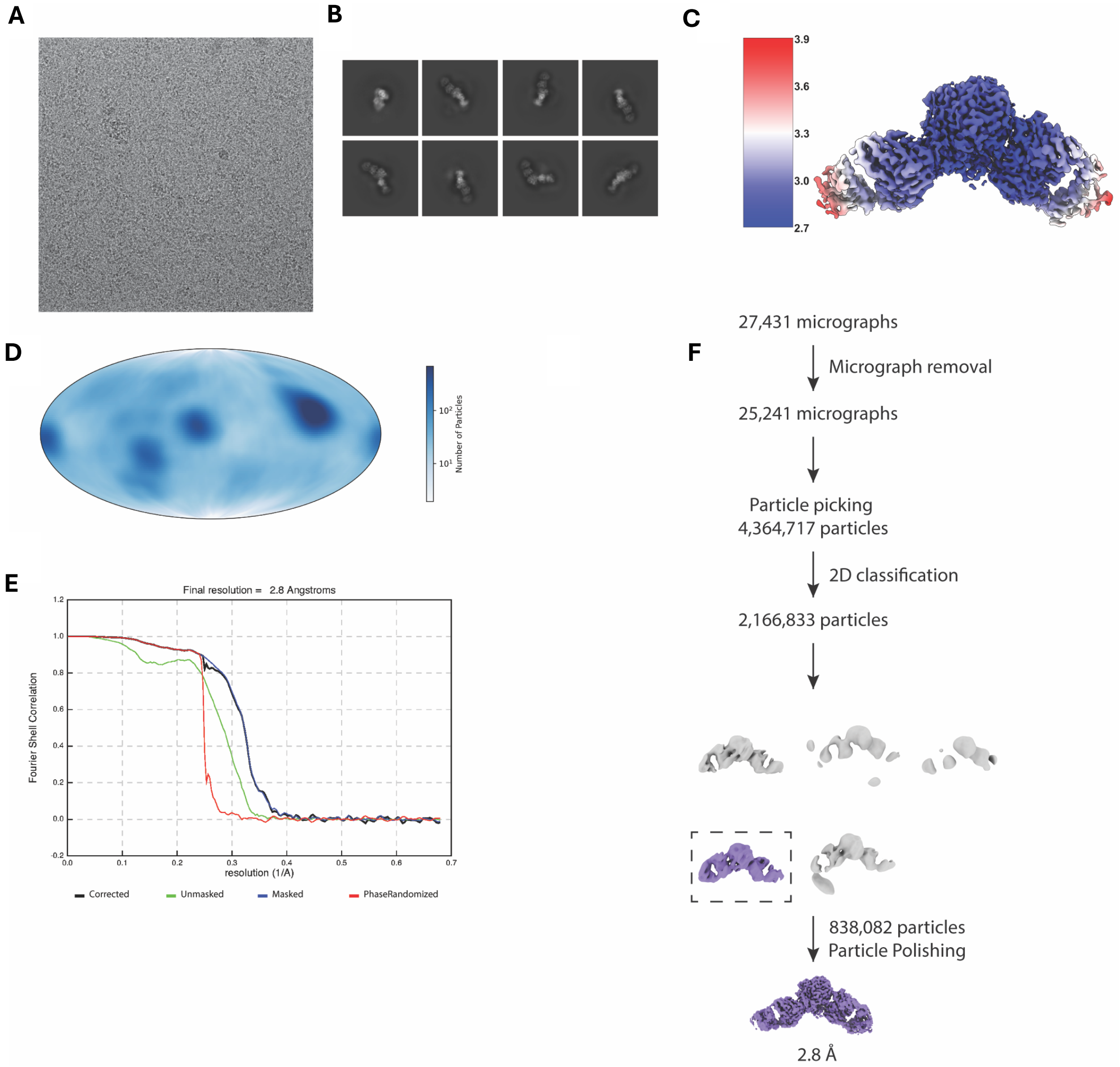

**fig. S10. Cryo-EM data collection. (A)** Representative micrograph of Dar1 and 1G01 Fabs bound to monomeric NA embedded in vitreous ice. **(B)** Selected 2D class averages of Dar1:1G01:NA complexes. **(C)** Reconstruction of complex filtered and colored by local resolution. **(D)** Angular distribution plot. **(E)** Fourier shell correlation (FSC) curves. **(F)** Schematic depicting the strategy for processing cryo-EM data.

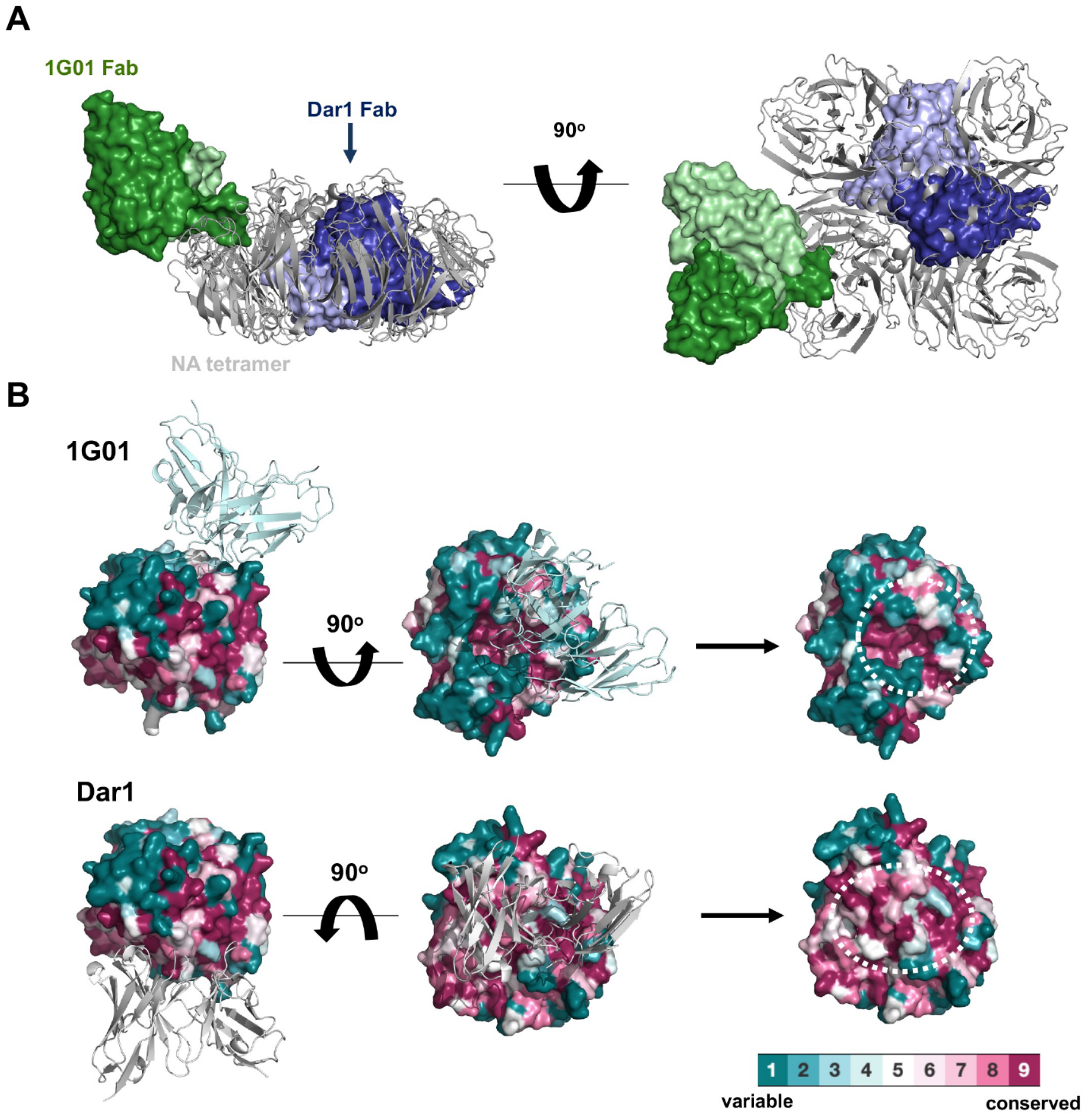

**fig. S11. Structural characterization of Dar1 and its epitope.** **(A)** Model of the NA tetramer with 1G01 and Dar1 Fabs docked highlighting that the latter recognizes an occluded interface when the tetramer is formed. The color scheme is used as in **Fig. 6C-D**. **(B)** 1G01 (top) and Dar1 (bottom) docked onto an NA monomer from J'57 N2 with conservation shown using ConSurf by analyzing 92 H2N2 and 7,576 H3N2 sequence-confirmed isolates from the Bacterial and Viral Bioinformatics Resource (same model as in **fig. S1B**). The general antigenic region recognized by each antibody is highlighted with a dashed white circle.

### table S1. Cryo-EM data collection, refinement and validation statistics

| | **Dar1:1G01:NA**<br>(EMDB-78520)<br>(PDB 37VO) |
|---|---|
| **Data collection and processing** | |
| Magnification | 165,000 |
| Voltage (kV) | 300 |
| Electron exposure (e–/Å$^2$) | 52.48 |
| Pixel size (Å) | 0.736 |
| Symmetry imposed | C1 |
| Initial particle images (no.) | 4,364,717 |
| Final particle images (no.) | 838082 |
| Map resolution (Å) | 2.8 |
| FSC threshold | 0.143 |
| Map resolution range (Å) | 2.7-4.0 |
| **Refinement** | |
| Initial model used (PDB code) | 3TIA, 6Q23 |
| Model composition | |
| Non-hydrogen atoms | 9,687 |
| Protein residues | 1,254 |
| Ligands | 1 |
| *B* factors (Å$^2$) (min/max/mean) | |
| Protein | 13.07/209.44/92.62 |
| Ligand | 85.01/103.52/93.02 |
| R.m.s. deviations | |
| Bond lengths (Å) | 0.005 |
| Bond angles (°) | 0.917 |
| Validation | |
| MolProbity score | 2.02 |
| Clashscore | 8.85 |
| Poor rotamers (%) | 1.56 |
| Ram-Z (RMSD) | |
| whole (N=1237) | -1.03 |
| helix (N=31) | -1.87 |
| sheet (N=500) | -0.14 |
| loop (N= 706) | -1.01 |
| Ramachandran plot | |
| Favored (%) | 94.17 |
| Allowed (%) | 5.51 |
| Disallowed (%) | 0.32 |
